# Sox9 marks limbal stem cells and is required for asymmetric cell fate switch in the corneal epithelium

**DOI:** 10.1101/2024.04.08.588195

**Authors:** Gabriella Rice, Olivia Farrelly, Sixia Huang, Paola Kuri, Ezra Curtis, Lisa Ohman, Ning Li, Christopher Lengner, Vivian Lee, Panteleimon Rompolas

## Abstract

Adult tissues with high cellular turnover require a balance between stem cell renewal and differentiation, yet the mechanisms underlying this equilibrium are unclear. The cornea exhibits a polarized lateral flow of progenitors from the peripheral stem cell niche to the center; attributed to differences in cellular fate. To identify genes that are critical for regulating the asymmetric fates of limbal stem cells and their transient amplified progeny in the central cornea, we utilized an in vivo cell cycle reporter to isolate proliferating basal cells across the anterior ocular surface epithelium and performed single-cell transcriptional analysis. This strategy greatly increased the resolution and revealed distinct basal cell identities with unique expression profiles of structural genes and transcription factors. We focused on Sox9; a transcription factor implicated in stem cell regulation across various organs. Sox9 was found to be differentially expressed between limbal stem cells and their progeny in the central corneal. Lineage tracing analysis confirmed that Sox9 marks long-lived limbal stem cells and conditional deletion led to abnormal differentiation and squamous metaplasia in the central cornea. These data suggest a requirement for Sox9 for the switch to asymmetric fate and commitment toward differentiation, as transient cells exit the limbal niche. By inhibiting terminal differentiation of corneal progenitors and forcing them into perpetual symmetric divisions, we replicated the Sox9 loss-of-function phenotype. Our findings reveal an essential role for Sox9 for the spatial regulation of asymmetric fate in the corneal epithelium that is required to sustain tissue homeostasis.

## Introduction

The corneal epithelium is a stratified, non-keratinizing tissue that serves as an external barrier to protect the eye against environmental insults, similar to the barrier function attributed to the epidermis in the skin. Like all stratified epithelia, the corneal epithelium undergoes constant cellular turnover. This turnover is the result of a differentiation trajectory, whereby cells exit the proliferative basal layer and terminally differentiate through the suprabasal and superficial layers of the tissue until they are eventually shed from the surface (desquamation) and lost in the tear film (Creamer et al., 1961; Haddad, 2000; Halprin, 1972; Potten et al., 1987). To match this constant cellular loss and maintain homeostasis, tissue renewal and replenishment is achieved by a population of resident stem cells that reside in the basal layer at the periphery of the tissue in a compartment known as the limbus (Cotsarelis et al., 1989; Schermer et al., 1986; Schlötzer-Schrehardt & Kruse, 2005).

In vivo lineage tracing analyses have demonstrated that long-lived corneal stem cells, termed limbal stem cells (LSCs) based on their location in the limbus at the periphery of the tissue, give rise to transient amplified cells (TACs), which exit the limbal niche and grow centripetally over several weeks before reaching the central cornea (Di Girolamo et al. 2015, Dora et al. 2015, Lobo, et al. 2016, Nasser et al. 2018, Park et al, 2019). Previously, we applied an intravital imaging technique, based on two-photon microscopy, to track the fates of individual cells throughout the corneal epithelium in the mouse eye. Quantitative clonal analysis of these data revealed the spatiotemporal organization of cell fates across the entire corneal epithelium (Farrelly et al. 2021). At the inner limbus, LSCs were found to undergo mostly symmetric cell divisions to expand their progeny. Subsequently, these short-lived transient amplified cells (TACs) divided in an increasingly asymmetric manner whereby their probability to terminally differentiate increased as they moved closer to the center of the cornea. Despite this continuous flux of progenitor cells in the basal layer, from the periphery to the central cornea, and their terminal differentiation from the basal to suprabasal/superficial layers, the tissue exhibits a remarkable ability to maintain stable cell numbers and overall tissue organization to support its barrier function. This phenomenon suggests a mechanism that instructs cells to undergo a specific mode of cell division (symmetric vs. asymmetric) depending on their precise location across the ocular epithelium, but the genes involved in such regulatory network remain elusive.

A major challenge in resolving the underlying mechanism of cellular fate is the dynamic nature of high turnover tissues, like the cornea, which is not sufficiently resolved by static experimental approaches. This is particularly important when studying the regulation of stem cell and progenitor populations that constitute only a minor fraction of the total cellular content in the tissue. Recent advancements in single-cell genomics have revealed a previously underappreciated heterogeneity among LSCs that may be critical for maintenance and repair of the ocular surface (Lee et al., 2022; Altshuler et al., 2021; Català et al., 2021; Collin et al., 2021; Dou et al., 2021; Kaplan et al., 2019; D. Q. Li et al., 2021; J. M. Li et al., 2021). These studies have not only challenged the conventional paradigm of a homogenous stem cell population but have also led to the identification of new specific markers, such as GPHA2, that distinguish between quiescent and actively dividing LSC subsets (Altshuler et al., 2021, Collin et al., 2021). This highlights the intricate regulatory mechanisms governing LSC activity, which can inform novel regenerative therapeutic strategies to treat diseases of the ocular surface. However, discrepancies in the expression of new, as well as established markers across the different studies underscore the complex interplay between LSCs, their progeny, and their microenvironment, necessitating higher resolution genomic approaches for clearer delineation.

Advanced in vivo imaging techniques and cell-tracking strategies are pivotal in this endeavor, as they allow for the direct observation of cellular dynamics. This capability is essential for distinguishing between symmetric and asymmetric cell fates and for assessing their respective contributions to tissue structure and function, as well as pathologies which are largely characterized by abnormal cell proliferation and/or differentiation. Here, building on recent advances in genomics and mouse genetics, we ventured to identify the genes and regulatory networks that control the asymmetry of cell fate in the ocular surface epithelium. We identified Sox9 as a key transcription factor that is preferentially expressed in LSCs and demonstrate its necessity for proper regulation of asymmetric cell fate, which is critical to maintain corneal epithelial homeostasis.

## Results

### A high-fidelity reporter for analyzing cell cycle dynamics in the live cornea

Since LSCs and their progenitors (TACs) represent only a fraction of cells that make up the epithelium, we sought to enrich for these populations to interrogate the mechanisms that regulate the mode of cell division and therefore their fate. To do this, we implemented a dynamic in vivo reporter mouse model (CycB1-GFP) that enabled us to visualize the activity of actively cycling cells in the corneas of live mice (**Fig. 1A**). CycB1-GFP transgenic animals express an enhanced green fluorescent protein (EGFP) that is fused to the N-terminus of the Cyclin B1 (Ccnb1) gene (Klochendler et al., 2012). Since the N-terminus is targeted for degradation in a cell-cycle dependent manner, only cells that are actively cycling (S-G2-M stages) express the CycB1-GFP protein.

**Figure 1.**
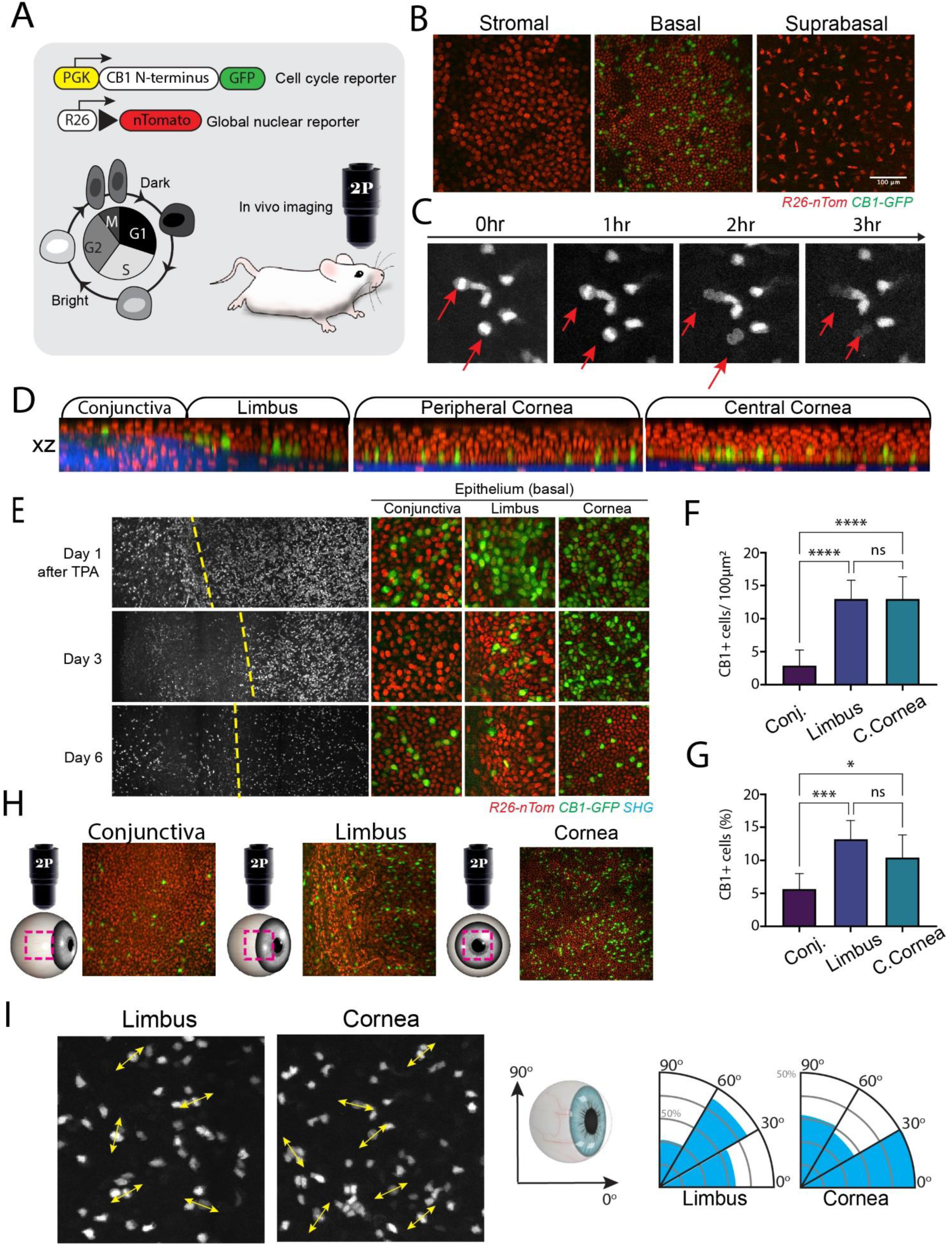
Capturing cell cycle activity in the live mouse cornea. (A) Genetic alleles of in vivo cell cycle reporter (Cyclin B1-GFP) used to sort stem and progenitor cells in the corneal epithelium. Cell cycle diagram displays how brightness of the GFP reporter changes throughout the cell cycle. (B) Individual planes from a series of optical sections of the mouse cornea taken at the indicated depths. (Also see <u>Movie S1</u>.) (C) Representative snapshots from time-lapse imaging of the cornea in a live CB1-GFP mouse. Only actively cycling cells are labelled with GFP (Also see <u>Movie S2</u>). (D) Reconstructed side view (XZ) of limbus and cornea from a CB1-GFP mouse based on 2-photon intravital imaging. (E) Panel shows representative snapshots from a time course live imaging experiment. The limbus and peripheral cornea were imaged at high magnification at the indicated times after treatment with TPA. Dotted line indicates the limbal-corneal border. (F-H) Optical sections of a CB1-GFP mouse eye imaged at the indicated epithelial compartments. The globally expressed nuclear tomato reporter allowed for visualization and quantification of total cell numbers. Graphs show quantification of the number of CB1-GFP+ cells (top panel) and percentage of CB1-GFP+ cells in each epithelial compartment (n = 6 [conjunctiva] or 8 [limbus + cornea] sampled images per epithelial compartment from 2 mice, one-way ANOVA). *p < 0.05, **p <0.01, ***p < 0.001, ****p < 0.0001. (**I**) Representative frames from time-resolved live imaging series, showing CycB1-GFP+ cells located in the basal layer of the limbus and central cornea. Arrowheads denote the axis of the daughter cell separation during mitosis, which are quantified and shown ass radial graphs (right panel) p = 0.86 (n = 35 cells [limbus] and 35 cells [limbus] sampled from 6 time-lapse datasets.

We performed time-lapse imaging in the corneas of live mice to validate reporter dynamics in actively cycling cells. We confirmed that only cells in the basal layer were GFP+ as only basal-layer cells in the corneal epithelium are capable of dividing (**Fig. 1B-D; <u>Movie S1</u>**). Using real-time imaging we visualized cells in the live cornea undergoing mitoses, which validated the fast kinetics of the in vivo cell cycle reporter (**Fig. 1C; <u>Movie S2</u>**). To further test the fidelity of the reporter, we topically administered 12-O-Tetradecanoylphorbol-13-acetate (TPA) to mouse corneas, which has a strong mitogenic effect on epithelial tissues (Taketani et al, 1983). Only one day after treatment, we saw massive increases in the number of GFP+ cells across the ocular epithelium, which was more pronounced in the limbus and central cornea (**Fig. 1E)**. One week after treatment, the number of GFP+ cells had declined and were comparable to untreated eyes. Taken together, we demonstrate here a powerful tool for visualizing cell proliferation dynamics in a living eye.

To capture possible imbalances in the rate of cell proliferation between the epithelial compartments, we performed image analysis and quantified the number of GFP+ cells in different areas of the corneal epithelium. Our data showed that actively cycling stem cells and progenitors are distributed uniformly within the basal layer of the limbus and cornea (**Fig. 1F-H**). This suggests that the rate of proliferation alone, does not impose a growth pressure between these compartments. To this point, we observed from our time-lapse imaging data from live corneas of CycB1-GFP mice that all cell divisions are planar with both daughter cells separating within the basal layer (**Fig. 1C, <u>Movie S2</u>**). This implies that the orientation of the mitotic spindle is likely not the mechanism by which corneal epithelial cells regulate their asymmetric fate and terminal differentiation. We then asked if the orientation of the cell divisions, in respect to the radial axis connecting the limbus to the center of the cornea, plays a role in the centripetal flux of progenitors within the basal layer. Our analysis showed that even though there were measurable biases for specific angles; on average, cell divisions in the limbus and cornea were oblique to the radial axis of the cornea (**Fig. 1I)**. From these data, we infer that the orientation of mitosis does not directly determine the fate of LSCs and TACs in the cornea.

### Single-cell transcriptomic analysis of actively cycling cells identifies key transcription factors in distinct progenitor populations in the corneal epithelium

To develop a greater understanding of the genes and molecular pathways that regulate the fate of LSCs and TACs, we harvested corneas from adult mice and performed Fluorescence Activated Cell Sorting (FACS) to purify only GFP+ corneal epithelial cells. (**Fig. 2A; Fig. S1A**). Cell filtering and unbiased clustering defined 8 cell populations (**Fig. 1B, C; Fig. S2A**). Direct comparison with published single-cell RNA sequencing datasets of mouse corneal epithelial cells showed an enrichment in proliferative basal cells (**Fig. S1B, C**). Moreover, we confirmed that the majority of cells in our dataset expressed basal cell markers, were actively cycling and were healthy, according to low expression of apoptotic markers (**Fig. S1D**). Taken together the initial analysis highlighted our ability to specifically interrogate gene expression in single LSCs and corneal TACs with high resolution.

**Figure 2.**
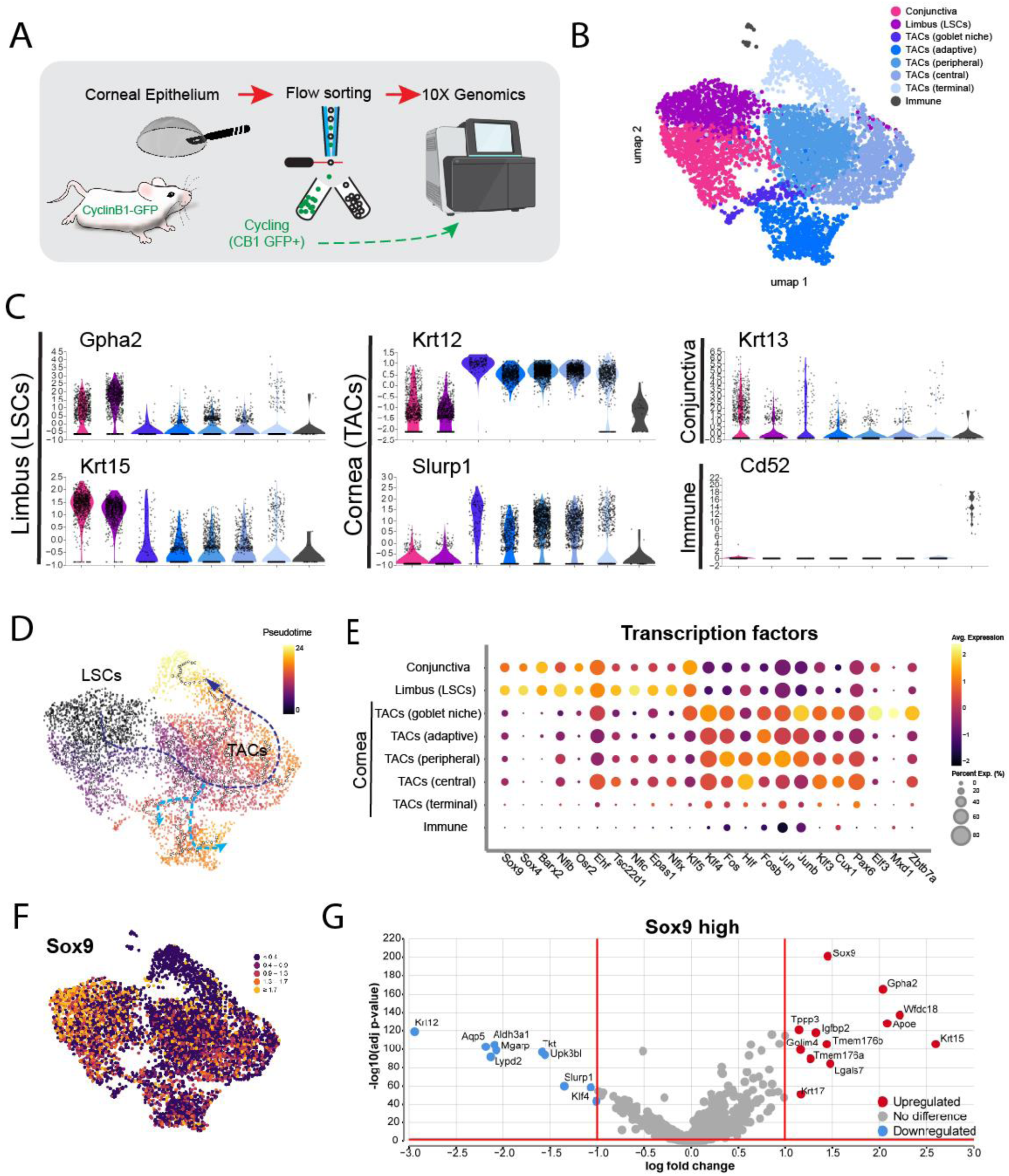
Single cell transcriptomic analysis of corneal stem and progenitor cells. (A) Experimental design for isolating and profiling actively cycling cells in the mouse ocular epithelium. (B) UMAP plot presentation of GFP+ cells sorted from corneas of CB1-GFP mice. Unbiased clustering of 5545 cells (2 pooled samples, 30 and 22 corneas total) revealed 8 distinct groups: Conjunctiva (860 cells), Limbus (LSCs; 823 cells), TAC (goblet niche; 150 cells), TACs (adaptive; 702 cells), TACs (peripheral; 1443 cells), TACs (central; 875), TACs (terminal; 668 cells), Immune (24 cells). (C) Stacked violin plots of differentially expressed markers used to define cluster identity. The y-axes represent expression level. (D) Inference trajectory analysis. Dotted lines show inferred pseudotime trajectories of corneal progenitors. (E) Dot plot of top differentially expressed transcription factors. (F) Umap plot for Sox9. (G) Volcano plot comparing differentially expressed genes between Sox9-high (>1.3 log) or low (<1 log) expressing cells.

Unique cell identities were classified based on gene expression, which revealed clusters of conjunctival (1), limbal (1), corneal (5), immune (1) cells. Conjunctival cells were primarily identified based on the expression of Keratin 13 (Krt13), Keratin 19 (Krt19) (Altshuler et al., 2021; Ramirex-Miranda et al. 2011, Kasper et al., 1988), as well as the lack or low expression of corneal and limbal markers Keratin 12 (Krt12) (Kasper et al., 1988; Kurpakus et al., 1994; Liu et al., 1993) and Glycoprotein Hormone Subunit Alpha 2 (Gpha2) (Altshuler et al., 2021), respectively (**Fig. 1C, Fig. S2B**). Similarly, the limbal cluster was defined based on Krt15 and Gpha2 expression (Altshuler et al., 2021; S. Yoshida et al., 2006) and low expression, of both conjunctival and corneal markers (**Fig. 1C, Fig. S2B**). The corneal-specific Keratin 12 (Krt12) was detectable in 5 distinct clusters (**Fig. 1C, Fig. S2B**).

The smallest of those corneal TACs clusters displayed hallmarks of the previously described compound corneal niches, such as expression of Krt12, Krt19 and Mucin 4 (**Fig. 1C, Fig. S2B**) (Pajoohesh et al., 2012). These compound niches were shown to give rise to goblet cells in the corneal epithelium that are distinct from those in the conjunctiva and play a role in the pathology of limbal stem cell deficiency. Other unique genes associated with the goblet niche population are Krt6b, Krt78, Mal, Nccrp1, Plac8, Psca and the transcription factor Hopx (**Fig S2B**). Another distinct cell cluster within the cornea supergroup was characterized by the expression of Urah, a gene that was previously described to be highly expressed in a group of distinct corneal progenitors proposed to play a role in “adaptive regeneration” (Lin et al, PNAS 2023) (**Fig S2B**). Similarly, this cluster expresses Fabp5, Sparc and Krt16, consistent with previous reports. In addition to Krt12, progenitor cells in all the corneal clusters express high levels of Slurp1, Mgarp and Aqp5 (**Fig. S2B**).

To investigate the relationships between our identified cell clusters and resolve the differentiation trajectory of cells that maintain the corneal epithelium, we performed trajectory inference analysis. A clear trajectory emanating from the limbal cluster progressed through the TAC-peripheral, TAC-central, and TAC-terminal clusters, suggesting that these represent consecutive stages of terminal differentiation of the TACs that emerge from the limbus (**Fig. 2D**). The trajectory from the LSC cluster also showed distinct branching points towards the “goblet niche” and “adaptive” TAC clusters. Next, we performed differential expression analysis of genes that encode for transcription factors to identify key transcriptional regulators of the distinct cellular identities found within the stem and progenitor cell pools of the corneal epithelium. Our data showed that LSCs express high levels of Sox9, Sox4, Barx2, Nfib1 and Osr2 (**Fig. 2E)**. Expression of these genes decreases as the TACs exit the niche, and in turn there is upregulation of Klf4, Fos, Hif, Jun, Junb, and Klf3. (**Fig. 2E)**. While the five corneal clusters showed similar expression for the same transcription factors that differed from the limbal cells, the “goblet niche” progenitors additionally expressed high levels of Elf3, Mxd1 and Zbtb7a. Expression of all these transcription factors waned in the TAC-terminal cluster, suggesting that these cells may be the ones closest to terminal differentiation as also indicated by the trajectory analysis.

### Identification of the transcription factor Sox9 as a putative limbal stem cell marker

To address our original quest for genes which regulate the switch from symmetric cell fate in LSCs to asymmetric in TACs, we identified Sox9 as one of the transcription factors that are differentially expressed between the limbal and the corneal clusters (**Fig 2E, F**). The transcription factor Sox9 has long been known for its role as a regulator of stem and progenitor cell activity in other ectodermal-derived appendages (Kadaja et al., 2014; Poché et al., 2008; Scott et al., 2010; Vidal et al., 2005). Given the many parallels in gene expression patterns between the cornea and the hair follicle, where Sox9 has an established role in regulating hair follicle stem cell fate, we sought to explore the expression of Sox9 within our single cell dataset We confirmed that Sox9-high cells express all the typical limbal markers including Gpha2 and Krt15 (**Fig. 2G)**. Other highly expressed genes within the Sox9-high population are Wfdc18, Tppp3, Tmem17a, Tmem17b, Igfbp2, and Lgals7 (**Fig. 2G)**. Although transcript levels are not necessarily indicative of functional activity, we predicted that Sox9 may be functionally more important in LSCs than in corneal epithelial cells.

To interrogate the role of Sox9 in homeostatic maintenance of the corneal epithelium, we utilized a two-photon-microscopy-based system to visualize whole eyes in live mice without compromising the structural integrity of the tissue (Farrelly et al., 2021). To validate Sox9 expression within the murine cornea *in vivo*, we imaged the corneas of adult *Sox9^IRES-EGFP^*(Sox9-EGFP) mice (Chan et al., 2011), which express EGFP from the endogenous Sox9 locus (**Fig. 3A, B**). In these animals, GFP expression is almost exclusively restricted to the limbal compartment. Moreover, Sox9-EGFP levels have been previously shown to correlate with endogenous Sox9 levels (Formeister et al., 2009). Here, we observe Sox9-EGFP is expressed at two steady-state levels whereby Sox9 activity is highest in LSCs residing in the niche and lowest in progenitors that have exited the niche (**Fig, 3B**).

**Figure 3.**
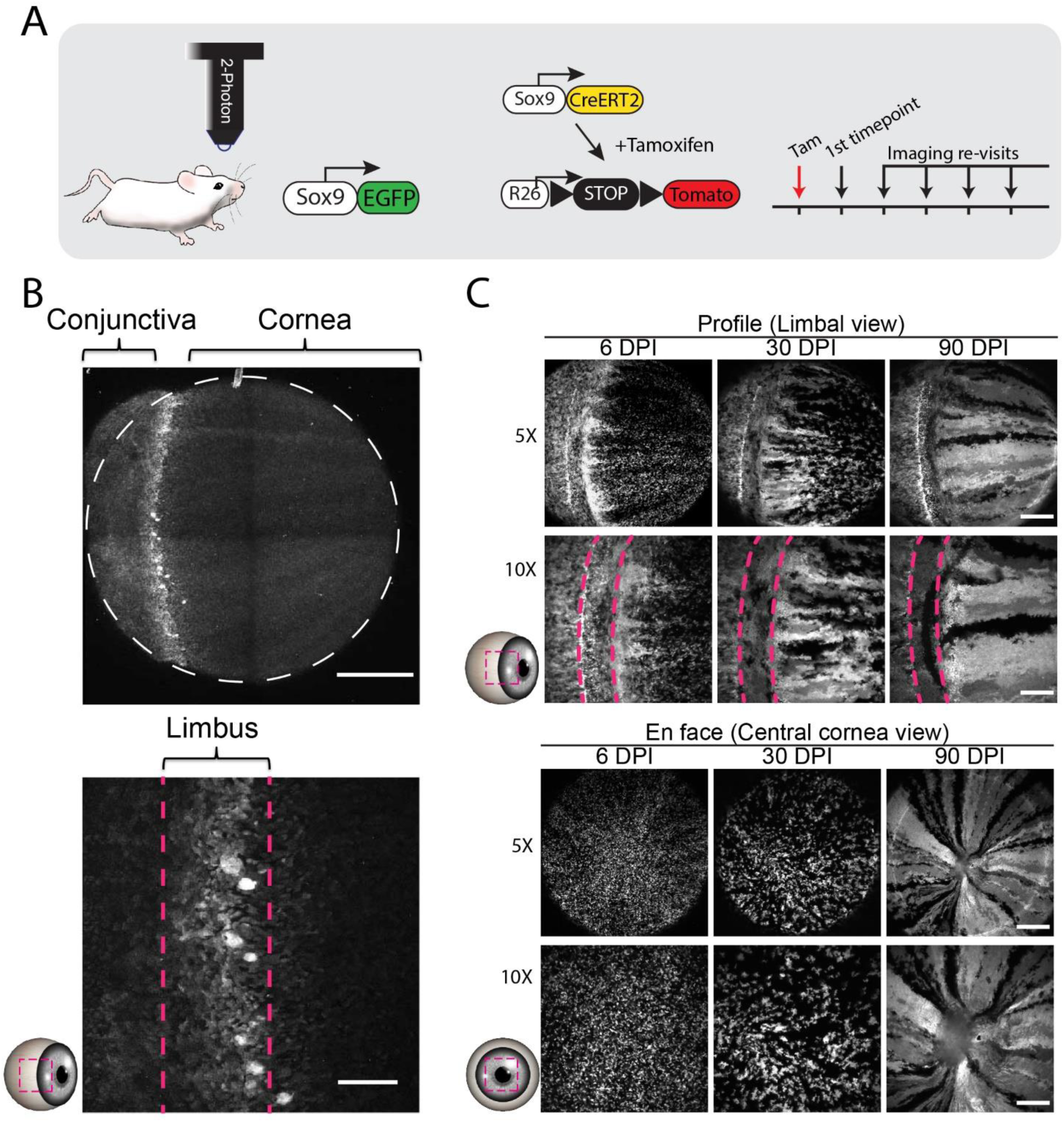
Sox9 marks long-lived limbal stem cells that maintain the cornea. (A) Genetic alleles for in vivo imaging. (B) Global and high-magnification views of the eye imaged at the indicated epithelial compartments show GFP expression predominantly at the limbus (n = 3 animals). Scale bars represent 500µm (global view) and 100µm (high magnification). (C) In vivo lineage tracing of Sox9CreER;tdTom cells by longitudinal live imaging. Representative tracing time series generated by reimaging the same eye at the indicated time points after tamoxifen administration (n = 5 mice). Lower panels show representative magnified views of the same compartment. Dotted lines indicate the margins of the limbus. Scale bars represent 500µm (panel B and 5x panels) and 250µm (panel C and10x panels). Abbreviations: DPI, Days Post Induction.

We next combined intravital imaging with an inducible Cre-recombinase-based *in vivo* lineage tracing strategy to resolve the activity and stemness potential of Sox9-expressing limbal cells in corneal homeostasis by longitudinal live imaging. These mice express a tamoxifen-inducible Cre recombinase from the endogenous Sox9 locus and a ROSA26LoxP-STOP-LoxP-tdTomato (R26-tdTom) Cre-reporter (Sox9CreERT2;tdTom) (**Fig. 3A**). To directly track the activity of Sox9CreERT2;tdTom lineage-traced cells, we acquired high resolution, full-thickness serial optical sections of the whole eye, including the limbus and central cornea, and then reimaged the same eyes longitudinally with the same acquisition parameters (**Fig. 3A, C**). By 1-month post-induction, we observed the emergence of long-lived lineages emanating from the limbus (**Fig. 3A**). Given that these lineages can only be sustained by LSCs (Amitai-Lange et al., 2015; Di Girolamo et al., 2015; Dorà et al., 2015), we concluded that Sox9 marks a population of LSCs as evidenced by their ability to contribute to long-term corneal homeostasis.

### Conditional deletion of Sox9 disrupts corneal epithelial homeostasis

To explore the tissue-specific role of Sox9 within the corneal epithelium, we implemented an inducible conditional gene targeting strategy (**Fig. 4A**). Sox9 floxed mice (Akiyama et al., 2002) were crossed to p63CreERT2 reporter animals (p63CreERT2;tdTom) to selectively delete Sox9 from basal-layer epithelial cells, the only cells that retain proliferative capacity and stemness potential (Cotsarelis et al., 1989), in adult corneas. By 2 months post induction (MPI), a proportion of Sox9 conditional knockout (cKO) corneas presented with hyperplastic growths at the central cornea, reminiscent of squamous metaplasia, and displayed extensive epithelial and stromal thickening at the limbus compared to control corneas (**Fig. 4B-E’’**). Compared to controls, Sox9-cKO corneas were thicker in the central cornea and limbal compartments (**Fig. 4H**). While stromal thickness was increased within the central cornea and limbal compartments of Sox9-cKO mice, changes in epithelial thickness were significant at the limbus, but not at the central cornea (**Fig. 4.4F-G**). This is likely due to the observed variance in plaque development/severity across Sox9-deleted animals. When Sox9-cKO corneas were scored on the severity of epithelial and/or stromal thickening, only about one-third of corneas (5/11 animals [7/21 corneas]) presented with moderate to severe epithelial thickening (**Fig. 4I-M**). This likely explains why epithelial thickening between experimental and control corneas is not significantly different (**Fig. 4F**). Based on these initial observations, we hypothesized Sox9 may play an important role in regulating the integrity of the corneal epithelium.

**Figure 4.**
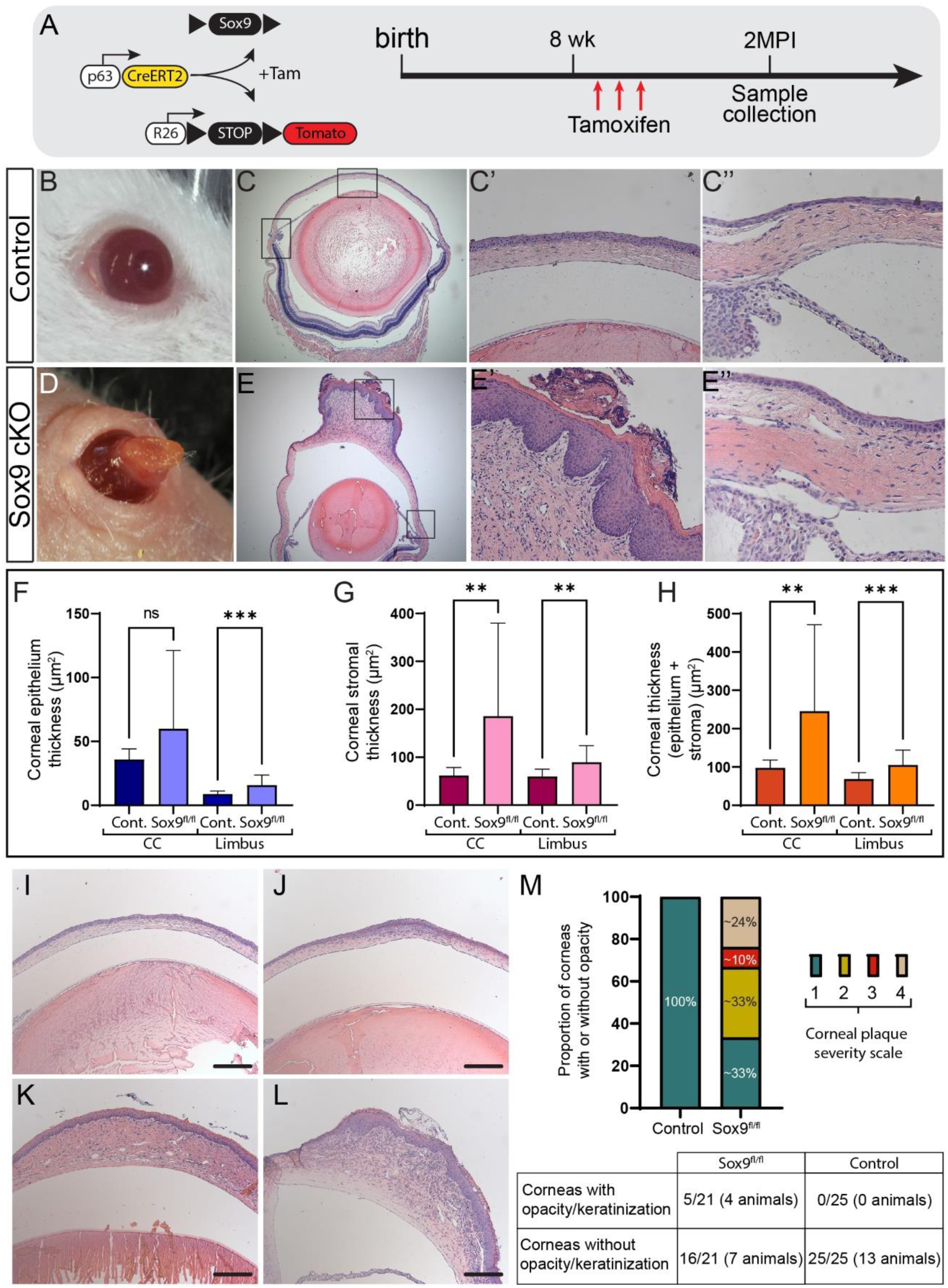
Sox9 cKO corneas develop squamous metaplasia. (A) Experimental timeline. Adult mice (p63CreER; Sox9 fl/fl cKO and control littermates) were treated with Tamoxifen daily for 3 days and analyzed 2 months post tamoxifen induction (MPI). (B-I) Gross morphology and representative histological sections of the central cornea (D, H) and limbus (E, I) of a control and Sox9-cKO cornea at 2MPI. (F-H) Quantifications of tissue thickness for the epithelium, stroma, and total corneal thickness in each epithelial compartment (control corneas: n = 13 animals [25 eyes]; Sox9-cKO corneas: n = 11 animals [21 eyes], Welch’s t test). (I-L) Representative histological sections depicting phenotype severity. Phenotype severity was scored from 1-4 as follows: 1) No gross morphological/molecular abnormalities (appear like controls), n = 7/21 corneas (6/11 animals); 2) Mild thickening of the epithelium and/or stroma, n = 7/21 corneas (6/11 animals); 3) Moderate stromal thickening with mild to moderate epithelial thickening, n = 2/21 corneas (1/11 animals); 4) Severe thickening of stroma, evidence of keratinization (loricrin expression), n = 5/21 corneas (4/11 animals). (M) Bar graph and table depicting distribution of corneas across phenotype severity categories and the proportion of corneas that developed opacities, respectively. Scale bars are 200 µm.

To begin characterizing our phenotype, we first validated the efficiency of our Cre/LoxP gene ablation system by performing immunostaining for Sox9 (**Fig. S3**). In control eyes, nuclear Sox9 staining was restricted to the basal layer cells within the limbus, whereas in the central cornea Sox9 faint staining could be observed in both the basal and suprabasal layers (**Fig. S3A/A’, C/C’**). In Sox9-cKO animals, Sox9 was not detected in either the limbus or central cornea, indicating efficient deletion of the Sox9 gene (**Fig. S3B/B’, D/D’**). Furthermore, we validated the epithelial specificity of our system by staining for Sox9 in retinal Müller glial cells, which are not of epithelial origin (**Fig. S3E/E’, F/F’**) (Poché et al., 2008).

Unlike control corneas, which exhibited a single K14+ basal layer, Sox9-cKO corneas displayed an expanded K14 layer in which K14 could be detected ectopically in the suprabasal and superficial layers of the epithelium (**Fig. S4A-D’**). Keratin 14 (Krt14) is a classic marker of basal-layer keratinocytes, the proliferative layer of the epithelium (Coulombe et al., 1993; Moll et al., 1982). Unlike the epidermis of the skin, the corneal epithelium is non-keratinized and mostly defined by the expression of Keratin 12 (Krt12) (Liu et al., 1993). In control animals, Krt12 expression can be detected throughout the suprabasal and superficial layers of the epithelium; however, it is not expressed within the limbus. In Sox9-cKO corneas, Krt12 is noticeably absent within the hyperplastic growths, while its expression can still be detected in the periphery of the corneal epithelium, outside the boundaries of the growths, where its expression is comparable to control eyes (**Fig. S4E-H’**). As Krt12 expression is unique to the corneal epithelium, its absence within the corneal plaques indicated a loss of corneal identity or cell fate switch.

Histologically, Sox9-cKO corneas displayed evidence of a keratinized surface, which had a striking resemblance to the stratum corneum of the epidermis (**Fig. S5**). Staining for Loricrin (Lor), a major structural protein of the epidermal cornified cell envelope, confirmed our suspicion that Sox9-cKO corneas were starting to take on an epidermal-like fate (**Fig. S5**). In Sox9-cKO corneas that displayed severe thickening and plaque growth (n = 5 corneas [4 mice]), Lor could be detected within the plaque growths, but not at the corneal periphery or limbus. This is consistent with the observation that K12 expression was properly patterned within Sox9-cKO corneas outside of the plaques. Collectively, these findings suggest Sox9 plays an important role in regulating corneal epithelial cell fate and identity as its loss resulted in a corneal-to-epidermal cell fate switch.

Within the corneal epithelium, cell fate decisions are largely influenced by the position of the cell within the tissue. At the limbus, LSCs divide symmetrically to maintain the stem cell pool but also to give rise to TAC progeny. As TACs exit the niche, they start to divide asymmetrically with increasing frequency giving rise to progenitors that will either terminally differentiate or divide into daughter cells with asymmetric fates where one terminally differentiates and the other is retained as a progenitor with proliferative capacity (Farrelly et al., 2021). Compared to controls, Sox9-cKO corneas with severe plaque development showed extensive hyperproliferation (**Fig. S6**). Staining for Ki67 revealed almost all basal cells, as well as cells in the suprabasal layers, were actively proliferating throughout the limbus and corneal epithelium compared to controls (**Fig. S6 A-D’**). We postulated that the increased proliferative activity throughout the corneal epithelium in the absence of Sox9 explains, at least in part, the observed thickening of the epithelium.

### Inhibition of terminal differentiation in corneal TACs phenocopies the loss of Sox9

To this end, we provided evidence that the transcription factor Sox9 is preferentially expressed in LSCs and that conditional ablation of Sox9 from the basal layer of the corneal epithelium leads to the development of squamous metaplasia at the central cornea. Although we hypothesized that loss of Sox9 would primarily affect the limbus, the most prominent phenotype in Sox9 cKO mice is the aberrant growths developing at the center of the cornea. Given that cell divisions switch from mostly symmetric in the limbus to increasingly asymmetric in the central cornea, with an increased propensity to terminally differentiate (Farrelly et al. 2021), we hypothesized that the transition from Sox9-high to Sox9-low expression as the TACs exit the limbal niche is required for their normal differentiation. Thus, we predicted that after conditional deletion of Sox9 TACs are unable to switch to asymmetric fates, without which they continue to divide symmetrically leading to cell crowding at the center of the cornea.

To test this hypothesis, we developed an inducible dual Cre and Tet system that would enable us to suppress differentiation (Differentiation-OFF) in single clones that we could visualize with a co-dependent H2BGFP reporter and track their behavior over time by live imaging (**Fig. 5**) (Farrelly et al., 2019). The premise of this genetic system is that active Notch signaling is required for the commitment to terminal differentiation of progenitors in stratified epithelia. Therefore, by enforcing total inhibition of canonical Notch, using the dominant negative MAML1 allele (Differentiation-OFF), we predicted that cells would be locked in a symmetric fate and undergo perpetual self-renewal. While H2BGFP labelled clones in control corneas could be observed differentiating normally and moving from the basal to the suprabasal layers of the epithelium over time, labelled clones in “Differentiation-OFF” corneas remained trapped in the basal layer (**Fig. 5B**). To further validate that suppressing asymmetric cell fate and differentiation of TACs leads to unchecked self-renewal, we performed quantitative clonal analysis. As expected, basal clones from normal corneas decayed over time, consistent with the short-lived nature of corneal TACs (**Fig. 5C**). In contrast, “Differentiation-OFF” clones continued to grow. While delamination of “Differentiation-OFF” cells was predictably suppressed for many days, it eventually started increasing after several days. This is likely due to crowding of cells which compete for space leading to cell extrusion from the basal layer (**Fig. 5B** see right panel, **<u>Movie S3</u>**).

**Figure 5.**
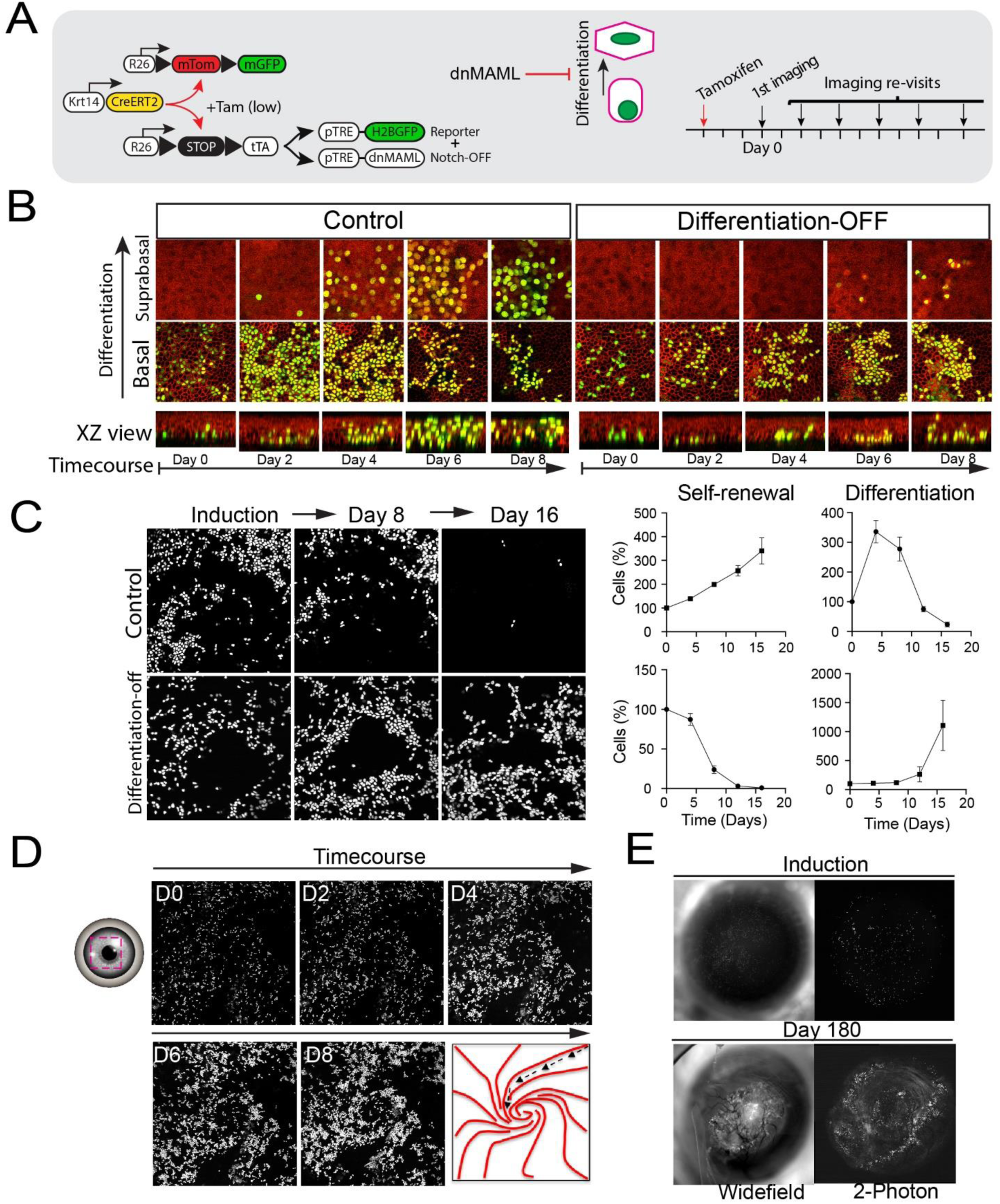
Cell migration and improper differentiation drive cellular crowding at the central cornea. (A) Allelic system and experimental strategy used to generate single, Differentiation-OFF clones in the basal layer of the corneal epithelium. Tamoxifen administration in low doses enables Cre-mediated excision of loxP-flanked sequences in a reporter allele and a tetracycline transactivator allele (tTA) in Keratin14-expressing cells. Membrane-localized GFP (mGFP) becomes expressed after excision of membrane-localized tdTomato (mTom) in single clones while excision of a STOP cassette in the ROSA26 locus allows tTA expression. tTA expression in single clones enables expression of two TetO alleles: a dominant negative MAML allele to suppress Notch signaling (Differentiation-OFF) and a reporter allele (TetO-H2BGFP) to label the single “Differentiation-OFF” clones with nuclear GFP. Control animals have only the one TetO-H2BGFP reporter allele. Labelled cells were followed over time by re-imaging the same areas of the cornea. (B) Labelled clones in control animals (n = 5) were tracked differentiating into the suprabasal and superficial layers of the cornea. Differentiation-OFF (n = 6 animals) clones were observed primarily in the basal layer with very few cells observed differentiating into the suprabasal and superficial layers. (C) Left panels show representative frames from a live imaging timecourse, showing the dynamics of control and Differentiation-OFF basal clones. Quantitative clonal analysis is shown in the graphs on the right. Two-way ANOVA, P<0.0001, n = 10 clonal fields tracked in 4 mice. (D) Representative frames from a live imaging timecourse of the central cornea from a Differentiation-OFF mouse. Over time, clones could be observed clonally expanding and migrating radially towards the central cornea. (E) Representative examples of corneas from Differentiation-OFF mice that developed squamous metaplasia in the central cornea.

To assess changes in migratory potential of “Differentiation-OFF” cells, we followed their fate by imaging the corneas of our “Differentiation-OFF" mice over time *in vivo*. Despite being unable to differentiate, “Differentiation-OFF” clones retained their ability to clonally expand and transit centripetally, leading to increasing crowding in the central cornea (**Fig. 5D**; **<u>Movie S4</u>**).

Moreover, several weeks after induction, we observed plaques developing in the central cornea that phenocopied those we previously saw in the Sox9-cKO mice. Altogether, these data provide support for a model whereby plaque growths in Sox9-cKO corneas are the result of TACs failure to switch to an asymmetric cell fate (cells trapped in basal cell identity), leading to their unchecked growth and crowding at the central cornea (**Fig. 6**). Therefore, Sox9 is a key transcription factor that regulates progenitor cell fate and is required to maintain corneal epithelial homeostasis.

**Figure 6.**
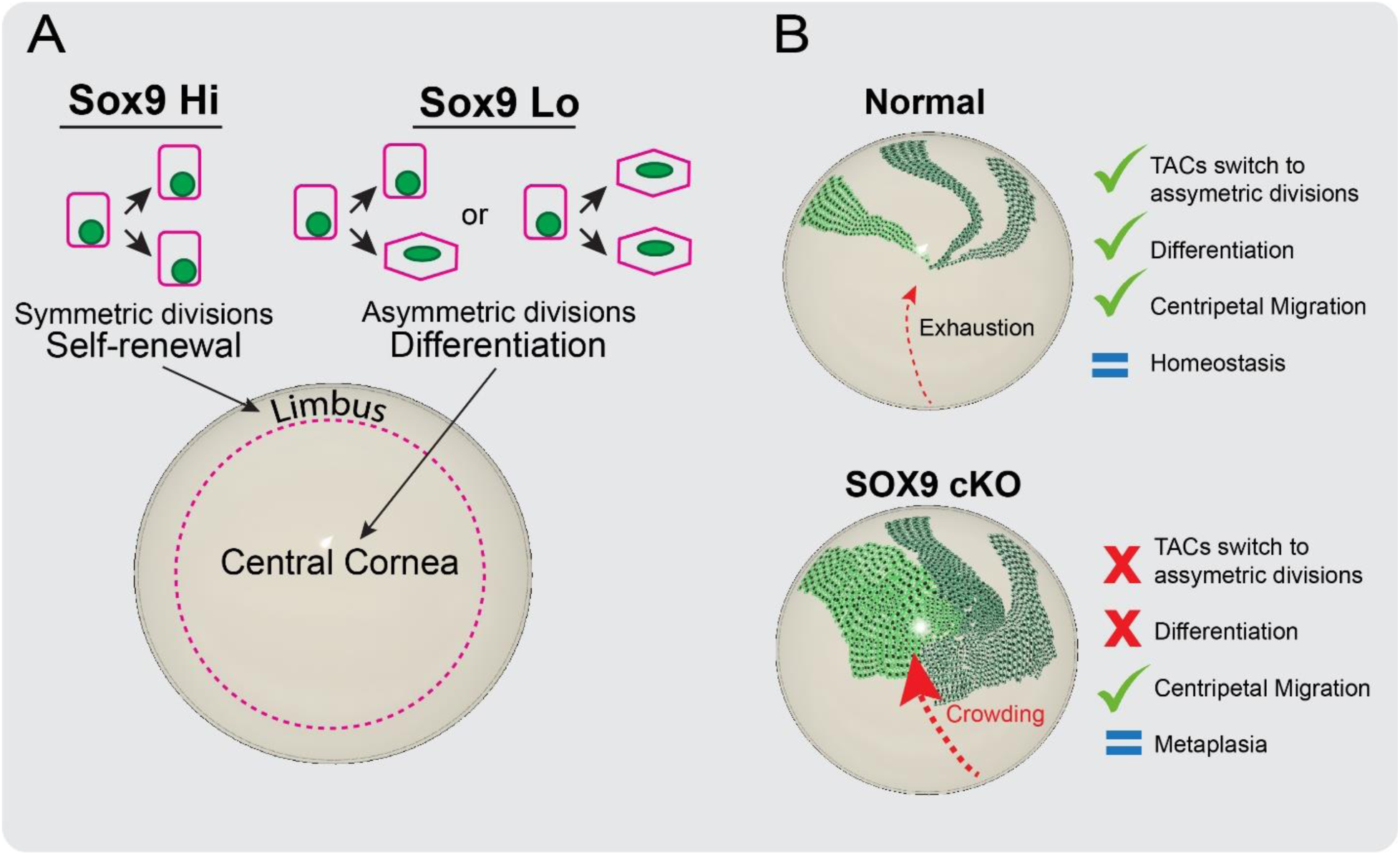
Mechanistic Model for the role of Sox9 in the regulation of stem cell fate in the ocular epithelium.

## Discussion

The corneal epithelium is a self-renewing tissue that is maintained by a population of tissue-resident stem cells (Cotsarelis et al., 1989). However, the molecular mechanisms which regulate the activity of LSCs and their progenitors are not fully known. The limbal niche is the specialized microenvironment in the limbus which regulates LSC function (H. S. Dua et al., 2005; Grieve et al., 2015; Tseng et al., 2016). As stem cells from the limbus divide symmetrically, they give rise to TACs, which migrate out from the limbal niche and transit centripetally towards the central cornea where they are eventually shed and washed away in the tear film (Lobo, et al. 2016, Nasser et al. 2018, Park et al, 2019, Farrelly, et al. 2021). This coordinated activity is achieved through a combination of cues from the niche which include niche factors like extracellular matrix, corneal nerve fibers, and coordinated signaling of various pathways (Gesteira et al., 2017; W. Li et al., 2007; Mei et al., 2012; Xu et al., 2023). Amongst these signaling pathways, several transcription factors have been shown to play a critical role in LSC maintenance and function (Bhattacharya et al., 2019; González et al., 2019; G. Li et al., 2015; Lu et al., 2012; Nakatsu et al., 2011; Sartaj et al., 2017; Shwartz et al., 2020; Vauclair et al., 2007). Our results in this study identified the transcription factor Sox9 as a putative marker of LSCs. Based on our single cell dataset and lineage tracing experiments, we determined that Sox9 is also expressed within the conjunctiva, a vascularized mucosal epithelium that is physically separated from the corneal epithelium via the limbus (Wei et al., 1996). However, it is well-established that the conjunctiva does not contribute to homeostatic maintenance of the corneal epithelium (S. H. Dua, 1998; Kruse et al., 1990; Shapiro et al., 1981; Wei et al., 1993, 1996). Thus, it is highly unlikely that any of the Sox9 lineage-traced corneal cells were of conjunctival origin.

The transcription factor Sox9 has been reported to be essential for cell proliferation and lineage commitment of adult human corneal epithelial stem cells in ex vivo cultures (Menzel-Severing et al., 2018). Sox9 is known to have a critical role in the maintenance of several adult tissues (Blache et al., 2004; Furuyama et al., 2011; Poché et al., 2008). Functionally, Sox9 has been described as a regulator of stem/progenitor cell maintenance and as having a role in cell fate specification (Aoki et al., 2003; Bastide et al., 2007; Kadaja et al., 2014; Passeron et al., 2007; Scott et al., 2010). However, the role that Sox9 plays in regulating corneal stem cell activity and fate in vivo was not previously addressed. By conditionally deleting of Sox9 in basal epithelial progenitors, we showed that the loss of Sox9 causes the corneal epithelium to become hyperproliferative, which suggests Sox9 primarily functions as a suppressor of symmetric cell division in the epithelium. Furthermore, the loss of Sox9 results in improper differentiation and keratinization of the corneal epithelium leading to the development of squamous metaplasia in the central cornea.

Previously, cKOs of Notch1 from the corneal epithelium were reported to develop a similar epidermal cell fate switch (Movahedan et al., 2013; Vauclair et al., 2007). In addition to the epidermal-like plaques, these animals had also displayed swollen eyelids. Thus, meibomian gland dysfunction was thought to be a contributing factor, at least indirectly, to the epidermal cell fate switch. Meibomian glands synthesize and secrete lipids that make up the protective tear film which covers the ocular surface under homeostatic conditions, (McCulley & Shine, 2003).

Disruptions to proper meibomian gland activity can lead to a disrupted tear film, making the corneal epithelium more susceptible to chronic irritation and injury (Chhadva et al., 2017). While meibomian glands in Notch1-deleted corneas developed into cyst-like structures and were devoid of meibum, the lipid secretion produced by the glands, meibomian gland dysfunction alone was not enough to produce the epidermal cell fate switch in Notch1-deficient corneas.

Although minor injuries are normally repaired by an LSC-mediated injury repair response, Notch1-deficient corneal cells could not properly repair the wounded epithelium and instead generated epidermal plaques at the central cornea (Vauclair et al., 2007). On the other hand, corneas that were protected from injury by having their eyelids stitched shut never developed epidermal growths on their eyes, providing further support for the role of Notch1 as a mediator of wound repair. Although we did not explore changes in meibomian gland physiology or wound repair in our Sox9-cKO corneas, it is not inconceivable to propose that the role of Sox9 in the cornea might also be one involved in injury repair rather than homeostatic maintenance. If this is the case, it would provide a reasonable explanation for as to why the epidermal growths in Sox9-cKO corneas vary so much in frequency and severity after 2 months. Perhaps the corneas that developed plaques had somehow been injured, resulting in a failed wound repair response and the growth of the epidermal plaques.

Cell fate and behavior is largely influenced by the cellular microenvironment. In most, but not all, self-renewing tissues, stem cells reside in a specialized microenvironment (niche) where they receive intrinsic and extrinsic signals to preserve their unique ability to self-renew and give rise to progenitor populations (Fuchs et al., 2004). Within the cornea, the activity and fate of LSCs and their progenitors is tightly regulated to ensure proper maintenance of the epithelium. As progenitors migrate out of the limbal niche, they embark on a differentiation trajectory that is influenced by their intrinsic differentiation potential and the cues they receive from the local microenvironment. For every cornea that developed epidermal growths, we were intrigued to see that these growths were always accompanied by thickening of the underlying stroma.

Previously, Vauclair et al., also showed that conditional ablation of Notch1 from the corneal epithelium resulted in stromal remodeling and the development of epidermal-like growths at the central cornea (Vauclair et al., 2007). While Vauclair et al., noted that the corneal-to-epidermal cell fate switch always developed after corneal injury, the authors acknowledged this cell fate switch likely involved cell non-autonomous signals mediated by the underlying stroma, which became largely vascularized because of increased FGF-2 signaling in the epithelium following Notch1 deletion. Thus, given the striking phenotypic similarities of Notch1-deleted corneas, it is plausible that the observed cell fate switch in our Sox9-deleted corneas may also be driven, in part, by cell non-autonomous signals from the underlying stroma.

## Acknowledgments

We thank Yana Kamberov, Andrew Vaughn, Patrick Seale and Aimee Payne for their valuable discussions and feedback that guided this study. We are incredibly grateful for Steve Prouty and the Penn Skin Biology and Diseases Resource-based Center for their technical support. We also acknowledge the support from the Institute for Regenerative Medicine and the entire stem cell community at Penn. P.R. was supported by a grant from NIH/NEI (R01EY030599) and HRFF grant from the State of PA. G.R. was supported by NIH/NIAMS pre-doctoral fellowship F31AR079899. O.F. was supported by NIH/NICHD training grant T32HD083185. Penn SBDRC was supported by center core NIH/NIAMS grant P30AR069589.

Author contributions: G.R. and P.R. conceptualized the study, designed the experiments, and wrote the manuscript. G.R. performed most experiments. L.O. and E.C. performed histological analyses. O.F., S.H., L.N. and C.L. performed the scRNAseq experiment. All authors discussed results and participated in the manuscript preparation and editing. P.R. supervised the study.

## Data and materials availability

Genomic data are available in GEO (GSE263151) Additional data sets and reagents presented in this study are available from the corresponding author upon request.

## Material and methods

### Mice

All procedures involving animal subjects were performed with the approval of the Institutional Animal Care and Use Committee (IACUC) of the University of Pennsylvania and were consistent with the guidelines set forth by the ARVO Statement for the Use of Animals in Ophthalmic and Vision Research. p63CreERT2 mice were created by J. Xu (Baylor College of Medicine) and obtained from A. Vaughan (University of Pennsylvania). K14CreERT mice were created by Valeri Vasioukhin (University of Chicago) and obtained from Elaine Fuchs (The Rockefeller University). R26loxp-stop-loxp-tdTom, R26loxp-stop-loxp-tTA, Ptre-H2BGFP, TetOS-dnMAML-GFP, and Sox9IRES-EGFP mice were obtained from The Jackson Laboratory. Sox9fl/fl mice come from The Jackson Laboratory and were obtained from V. Lefebvre (University of Pennsylvania). All mice used in this study were bred for multiple generations into a Crl:CD1(ICR) mixed background. Adult mice (8 weeks old) were used for experiments. Mice were housed in a temperature and light-controlled environment and received food and water ad libitum. Up to 5 mice of the same sex and similar age were housed in a cage. Mice were provided Bed-o’Cobs (The Andersons Lab Bedding), a porous cob material, as bedding and Shred-n’Rich nestlets (The Andersons Lab Bedding) for nesting and enrichment.

### Tamoxifen induction

To induce Cre-recombinase activation, mice were injected intraperitoneally (IP) with Tamoxifen (Sigma) dissolved in corn oil (Sigma). For lineage tracing experiments, Sox9-CreERT2;R26-tdTom mice were administered a single dose of 2 mg Tamoxifen. For lineage tracing experiments in Notch-suppressed K14Cre-ERT;R26-tTA;TetOS-dnMAML-GFP and control K14Cre-ERT;R26-tTA;pTRE-H2BGFP mice, a single dose of 0.2 mg Tamoxifen was administered. To induce floxed deletion of Sox9, p63-CreERT2;Sox9 fl/fl;R26-tdTom mice and littermate controls (Sox9 fl/fl;R26-tdTom) were administered a single dose of 2 mg Tamoxifen daily for 3 days. Subsequent experiments were conducted at the indicated times after induction.

### Intravital imaging of the mouse eye

Preparation of the mice for intravital imaging of the eye was performed with the following amendments to the previously described protocol (Rompolas et al., 2016). Mice were initially anesthetized with IP injection of ketamine/xylazine cocktail (0.1 mL / 20 g bodyweight: 87.5 mg / kg Ketamine, 12.5 mg / kg Xylazine). A deep plane of anesthesia was verified by checking pedal reflexes. The mouse head was stabilized with a custom-made stereotaxic apparatus that includes palate bar and nose clamp but no ear bars. Precision, 3-axis micro-manipulators are used to adjust the head tilt so that the eye to be imaged is facing up. A drop of eye gel (0.3% Hypromellose) was used as an optically neutral interface between the eye and a glass coverslip, and to prevent dryness and irritation to the tissue during the anesthesia and imaging procedure. After preparation and mounting is complete, the stage is placed on the microscope platform under the objective lens. A heating pad is used to keep a stable body temperature and vaporized isoflurane is delivered through a nose cone to maintain anesthesia for the duration of the imaging process. After each imaging session, the eyes were rinsed with PBS and the mice were monitored and allowed to recover in a warm chamber before being returned to their housing facility.

### Imaging equipment and acquisition settings

Image acquisition was performed with an upright Olympus FV1200MPE microscope, equipped with a Chameleon Vision II Ti:Sapphire laser. The laser beam was focused through 5X, 10X, 20X or 25X objective lenses (ZEISS FLUAR 440125-0000-000, N.A. 0.25; Olympus UPLSAPO10X2, N.A. 0.40; Olympus UPLSAPO20X, N.A. 0.75; Olympus XLPLN25XWMP2, N.A. 1.05). Emitted fluorescence was collected by two multi-alkali and two gallium arsenide phosphide (GaAsP) non-descanned detectors (NDD). The following wavelengths were collected by each detector: NDD1 419-458 nm; NDD1 458–495 nm; GaAsP-NDD1 495–540 nm; GaAsP-NDD2 575–630 nm. GFP and Tomato reporters were excited at 930 nm and their signal was collected by GaAsP-NDD1 and GaAsP-NDD2, respectively. Second harmonic generation (SHG) signal was generated using 850 nm or 930 nm excitation wavelengths and detected by NDD1 or NDD2, respectively. Serial optical sections were acquired in 10 µm steps, starting from the surface of the eye and capturing the entire thickness of the cornea (epithelium ∼40 µm, stroma/endothelium ∼80 µm). Expanded views of the cornea and limbus were obtained by acquiring a grid of sequential optical fields-of-view that were automatically stitched into one high-resolution tiled image using the microscope manufacturer software. Multi-day tracing experiments were done by re-imaging the same field of view or the entire eye at the indicated times after the initial acquisition. For each time point, inherent landmarks within the cornea, including the organization of the vasculature and collagen fibers (SHG), were used to consistently identify the limbus and navigate back to the original regions. Macroscopic images of the mouse eye were acquired under brightfield and fluorescence with an Olympus MVX10 Fluorescent Macro Zoom microscope fitted with Hamamatsu Orca CCD camera for digital imaging.

### Histology

Following euthanasia by CO2 narcosis and cervical dislocation in accordance with the American Veterinary Medical Association guidelines, whole globes were dissected and directly fixed in 1.3% paraformaldehyde in PBS for 1 hour at room temperature. Whole globes were washed in 1X PBS 2x10 minutes, placed in fresh 1X PBS, and stored at 4°C until further processing. For section staining, fixed globes were submitted to the Penn Skin Biology and Diseases Resource-based Center for paraffin embedding and sectioning. Slides were heated for 5-10 minutes at 70°C, deparaffinized using Everclear xylene substitute (2 x 5 minutes), and then rehydrated using a graded ethanol series (95% EtOH 2 x 2 minutes, 70% EtOH for 2 minutes, ddH2O 2 x 2 minutes). For antigen retrieval, slides were places in a 1:100 dilution of Antigen Unmasking Solution (Vector Laboratories) and heated in a boiling bath for 20 minutes and then left to sit at room temperature until solution cooled. Sections were washed as follows: ddH2O 1 x 5 minutes, 1 X PBS 2 x 3 minutes, 0.1% Triton X-100 in 1X PBS 2 x 3 minutes. Sections were incubated in blocking solution (10% normal goat serum, 0.1% Triton X-100 in 1X PBS) for 1 hour at room temperature and then incubated with primary antibody diluted in staining solution (2% normal goat serum 0.1% Triton X-100 PBS) overnight at 4°C. After 4 x 5-minute washes in 0.1% Triton X-100 in 1X PBS, tissue sections were incubated with secondary antibody and DAPI diluted in staining solution for 1 hour at room temperature. Secondary antibody incubation was followed by 4 x 5-minute washes in 0.1% Triton X-100 in 1X PBS and slide mounting using Fluor-Gel mounting medium (Electron Microscopy Sciences).

The following primary antibodies were used: Anti-KI67 (1:100, ab16667, Abcam), Anti-Sox9 (1:500, ab185966, Abcam), Anti-KERATIN 17 (1:300, 4543, Cell Signaling Technology), anti-hair cortex cytokeratin/K40 [AE13] (1:50, ab16113, Abcam), and anti-RFP (1:500, Rockland Immunochemicals, Cat # 600-401-379).

The following secondary antibodies were used: Alexa Fluor 488 Goat anti-Mouse (1:500, A11001, ThermoFisher Scientific), Alexa Fluor 488 Goat anti-Rabbit (1:500, A11008, ThermoFisher Scientific), Alexa Fluor 594 Goat anti-Rabbit (1:500, A11012, ThermoFisher Scientific), Alexa Fluor 594 Goat anti-Mouse (1:500, A11005, ThermoFisher Scientific), and DAPI (1:1000, 40043, Biotium).

### Image acquisition of stained tissue sections

Images of stained tissue sections were acquired using an Olympus BX51 equipped with a Hamamatsu Orca CCD camera or a Leica DM6 B equipped with a Leica DFC9000 GT fluorescent camera and a Leica DMC2900 brightfield camera.

### Corneal epithelial cell isolation for flow cytometry and sorting

Whole globes were isolated and the corneas, including the limbus and marginal conjunctiva, were dissected and placed into 2.4 units/µl of Dispase prepared in Keratinocyte Serum Free Medium and incubated at 37°C for one hour (sample 1) or overnight at 4°C (sample 2). For each cornea, the epithelium was gently peeled off, washed in 1 X PBS, and incubated in Trypsin-EDTA (0.25%) at 37°C for 20 minutes. Following incubation, tissue was processed into a cell suspension via successive rounds of manual pipetting with a P1000 tip. 2 ml of FBS and 6 ml of 1 X PBS were added to the cell suspension and cells were spun down at 1000-1500 rpm for 5 minutes. Following centrifugation, the supernatant was aspirated, cells were resuspended in 1X PBS, and the entire volume was filtered through a 40 µm cell strainer to obtain a single cell suspension. For cell staining, DAPI (1:1000) was added directly to the cell suspension and incubated for 10 minutes on ice. Following incubation, cells were spun down at 1000-1500 rpm for 5 minutes, supernatant was aspirated, and cells were resuspended for sorting in 2% FBS 5 mM EDTA in 1X PBS. Flow cytometry was performed on a BD LSRII cytometer (BD Biosciences), sorting on a BD FACS Aria II sorter (BD Biosciences). Only GFP+ live cells were collected from sorting. Flow cytometry data was collected and exported using BD FACs Diva software (BD Biosciences) and analyzed and plotted using FlowJo software.

### Library preparation and data generation

The FACS-generated cell suspension was used to generate a single-cell RNA-seq library according to the 10x Genomics protocol (Chromium Single Cell 3’ Library & Gel Bead Kit v3, PN-1000075). Single cell separation was performed using the Chromium Single Cell B Chip Kit (PN-1000074). The library was sequenced on an Illumina NextSeq500 instrument with a high-output mode (Illumina, 20024907) 150bp paired-end reading strategy according to 10X recommendations.

### Processing of 10X Genomics data

Cell Ranger (version 3.1.0) was used for primary data analysis (alignment, filtering, barcode and UMI counting) to determine transcript counts per cell (producing a gene-barcode matrix). Transcripts were mapped to the mouse reference genome, GRCm38 (refdata-cellranger-mm10–2.1.0/GRCm38.84)

### Downstream analysis of single-cell RNA sequencing data

The single cell RNA-seq dataset was processed, explored and visualised using Cellenics community instance (https://scp.biomage.net/) that is hosted by Biomage (https://biomage.net/) Filters for mitochondrial content (>7%), number of genes vs UMIs (linear regression, p-value=0.0003250) and doublet (scDblFinder, probability thresholding=0.56) were applied to exclude low quality cells. Data integration was performed using Harmony (2000 highly variable genes, LogNormalized). Dimensionality reduction was performed using 30 principal components with 87.09% variation explained, and normalized excluding ribosomal, mitochondrial and cell cycle genes. UMAP embedding was carried out using Euclidean distance metric and 0.6 minimal distance. Clustering was performed using Louvain at 0.3 resolution.

### Quantitative image analysis of CB1-GFP+ cells

Raw digital files from 2-photon imaging were acquired and saved in the OIB format using the microscope manufacturer’s software (FluoView, Olympus USA). Extended fields-of-view that encompass the entire ocular surface epithelium, including the cornea, limbus, and conjunctiva, were captured using a tiling method to reconstruct a single image from multiple full-thickness serial optical sections (FluoView, Olympus USA). To image the entire eye with the 10X objective lens, a matrix area of 2 ✕ 2 (XY) field-of-view with 10% overlap between tiles is defined. Using the motorized platform, the microscope automatically acquires the four fields-of-view in a sequential pattern and uses information from the overlapping margins to stitch the individual fields-of-view into a single image. Raw image files were imported into ImageJ/Fiji (NIH) using Bio-Formats for further analysis. For cell counts, supervised image segmentation and blob detection was performed on individual optical sections. Identified blobs were manually validated and their number, size, and signal intensity as mean gray values were measured. Images shown in figures typically represent maximum projections or single optical sections selected from the z stacks unless otherwise specified.

### Quantitative image analysis of corneal epithelium thickness

Images at the central cornea and limbus of stained H&E tissue sections were acquired using an Olympus BX51 equipped with a Hamamatsu Orca CCD camera or a Leica DM6 B equipped with a Leica DFC9000 GT fluorescent camera and a Leica DMC2900 brightfield camera. Length measurements were recorded using the straight line and Set Measurements functions in ImageJ/FIJI. Epithelial and stromal compartments were measured separately and then combined to generate values for total corneal thickness.

### Statistical analysis

Sample sizes were not pre-determined. Data were collected and quantified randomly, and their distribution was assumed normal, but this was not formally tested. Lineage tracing experiments were successfully reproduced under similar conditions using different mouse cohorts. The values of ‘‘n’’ (sample size) refer to number of corneal samples, unless otherwise indicated in the figure legends. Statistical calculations and graphical representation of the data were performed using the Prism software package (GraphPad). Data are expressed as percentages or mean ± SEM and two-tailed Student’s t test was used to analyze datasets with two groups, unless otherwise stated in the figure legends. For all analyses, p-values < 0.05 were designated as significant and symbolized in figure plots as *p < 0.05, **p < 0.01, ***p < 0.001, ****p < 0.0001, with precise values supplied in figure legends. No data were excluded from the analyses.

## Key Resources

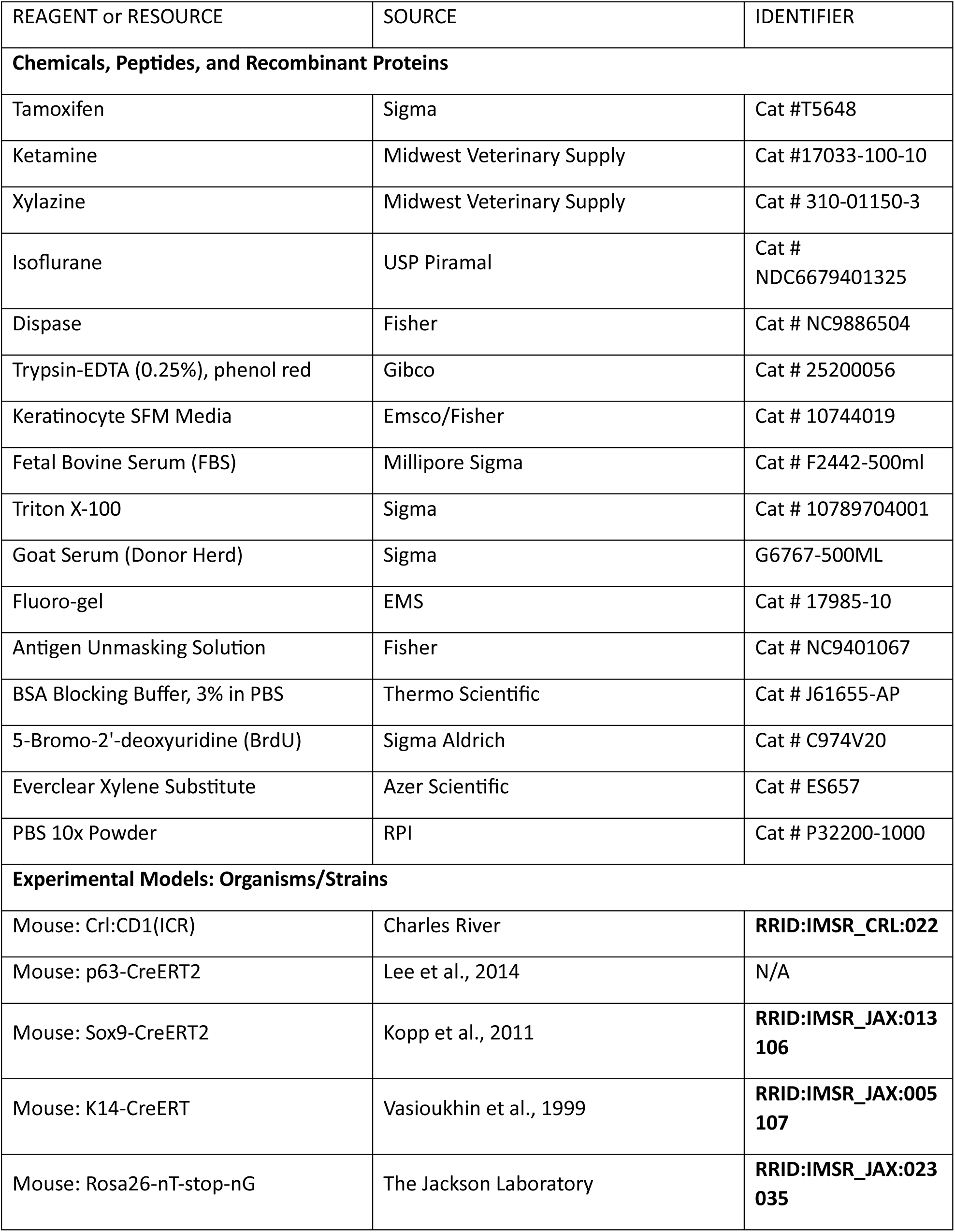
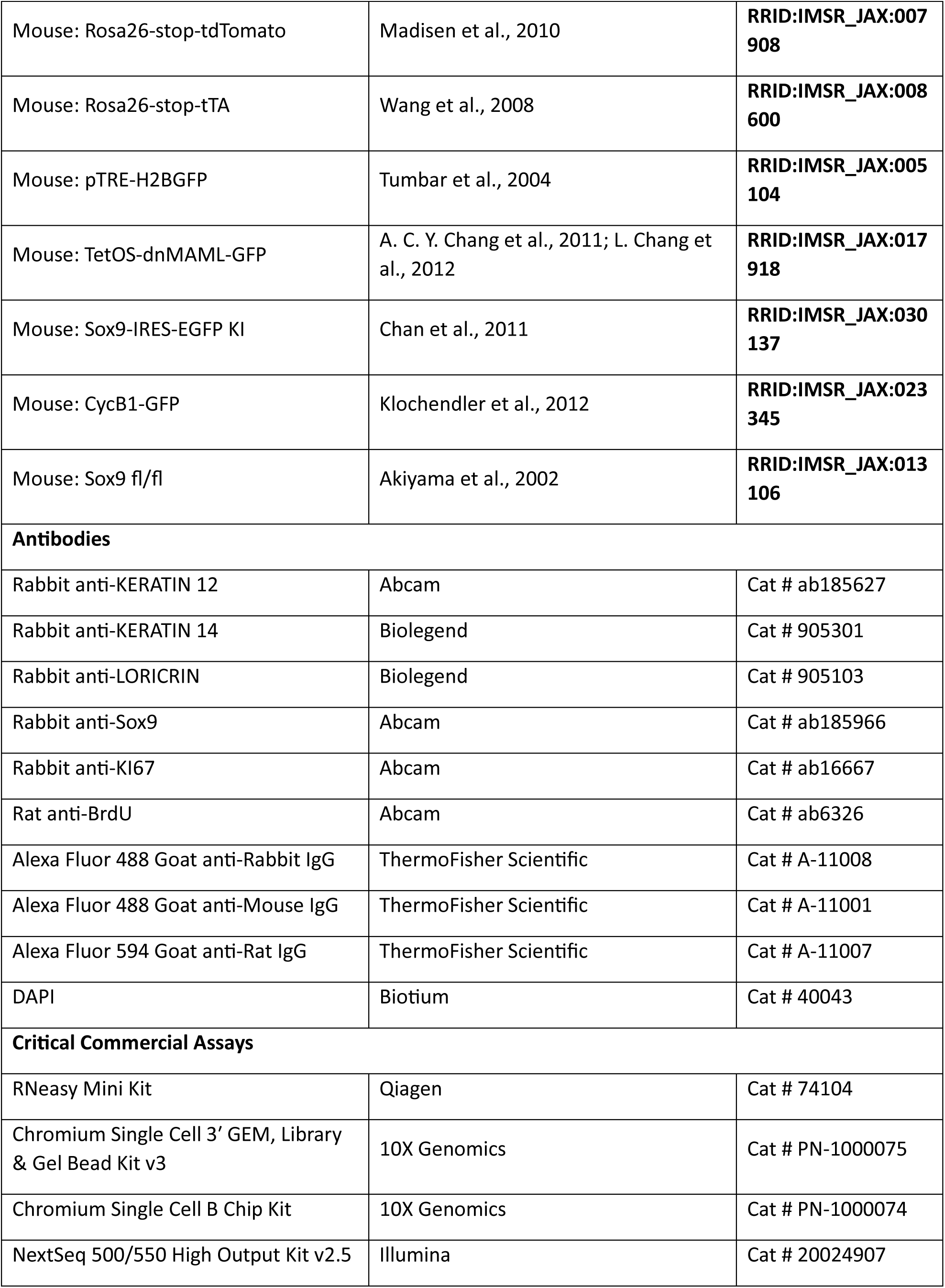
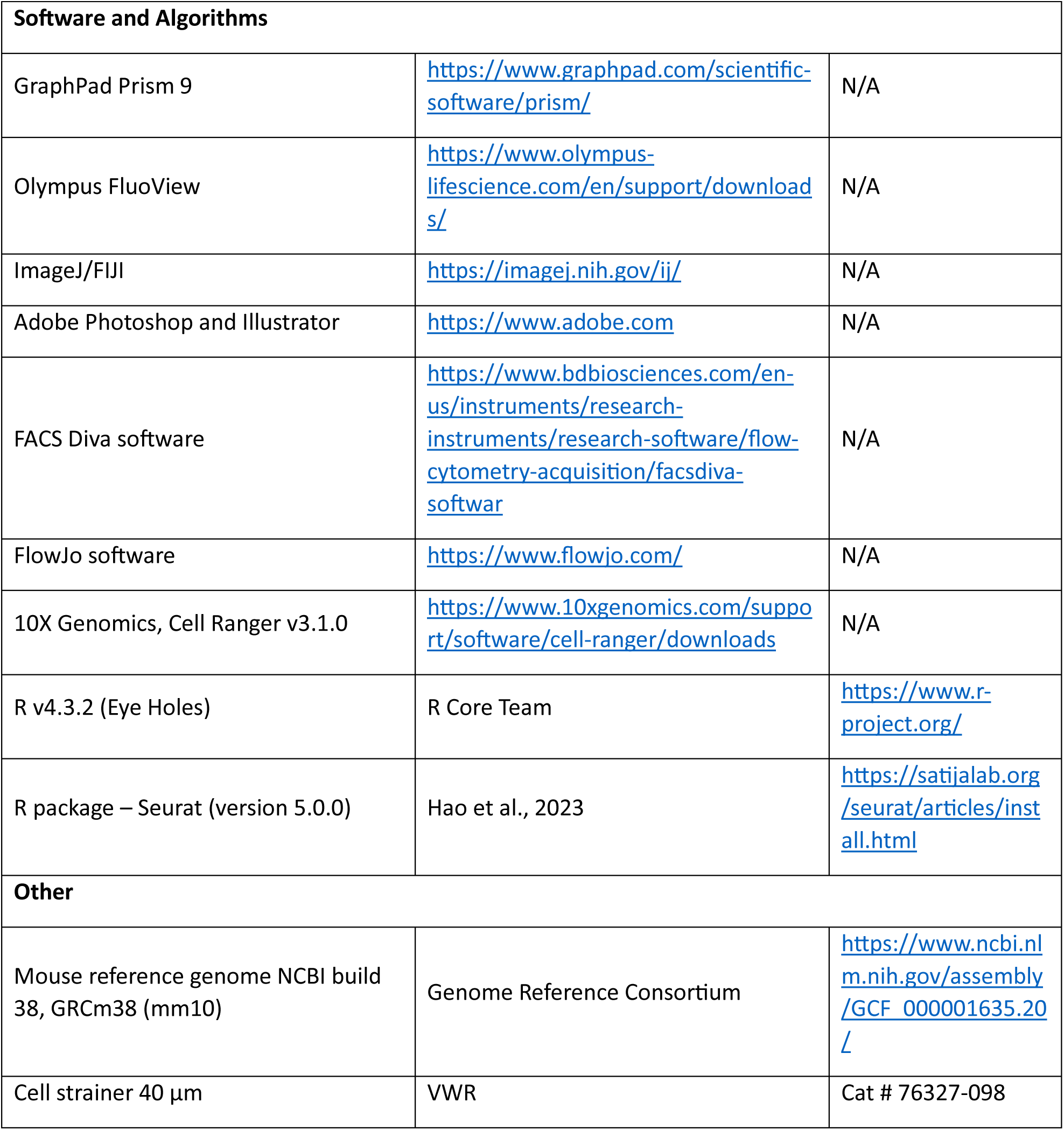

---

**Supplementary Figure 1.**
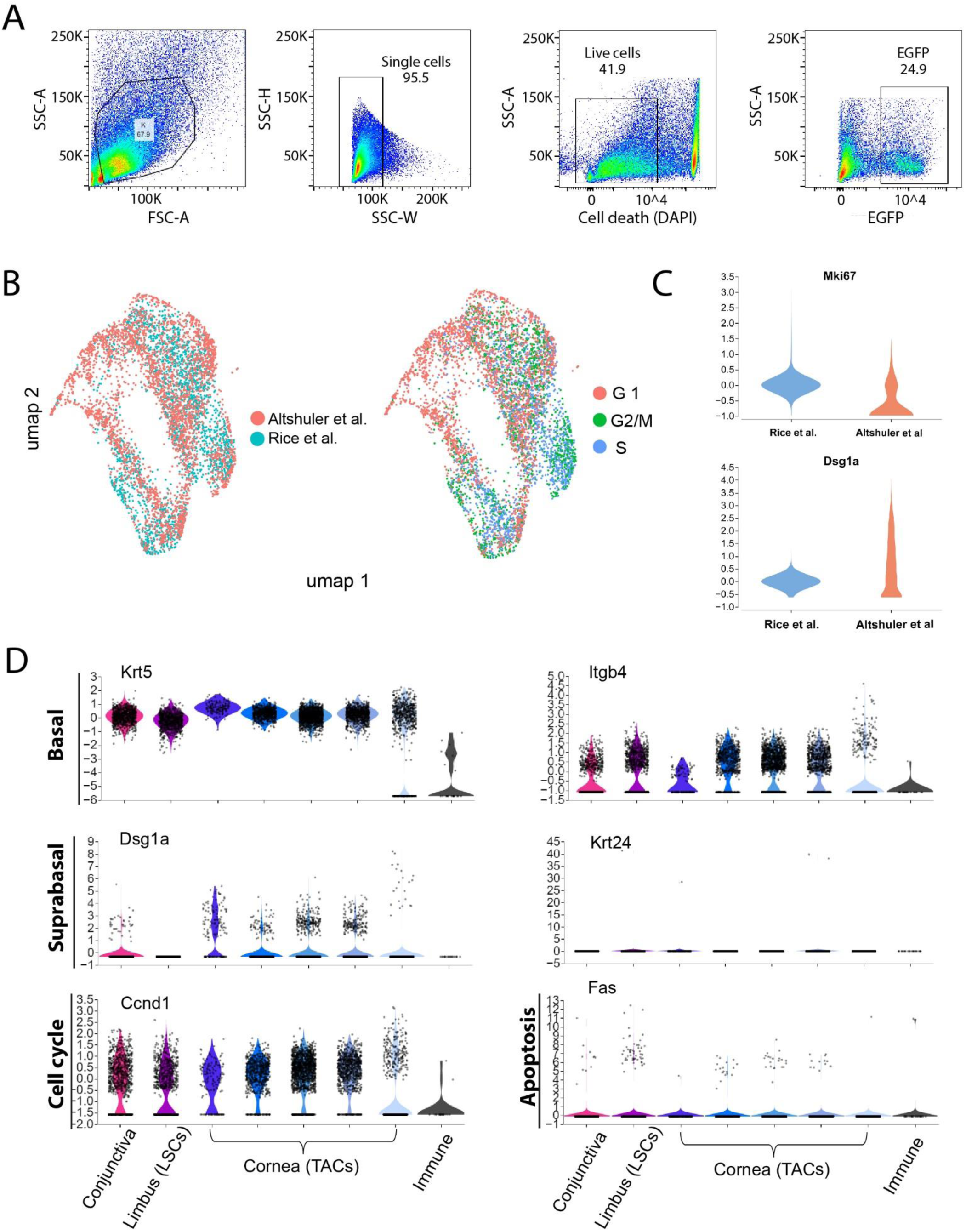
Single cell sequencing reveals CycB1-GFP sorted cells from the corneal epithelium have a mitotic, basal cell signature. (A) Representative images of FACS gating strategy used to sort for GFP+ cells from Cyclin B1-GFP animals. (B) Umap plots of integrated single cell datasets comparing this study with a published study of single cell transcriptomic analysis of bulk corneal epithelial cells (*Altshuler et al. 2021*). (C) Violin plots of proliferation (Mki67) and differentiation (Dsg1a) markers. (D) Violin plots of basal, suprabasal, cell cycle and apoptotic markers of mouse corneal epithelial cells profiled in the present study.

**Supplementary Figure 2.**
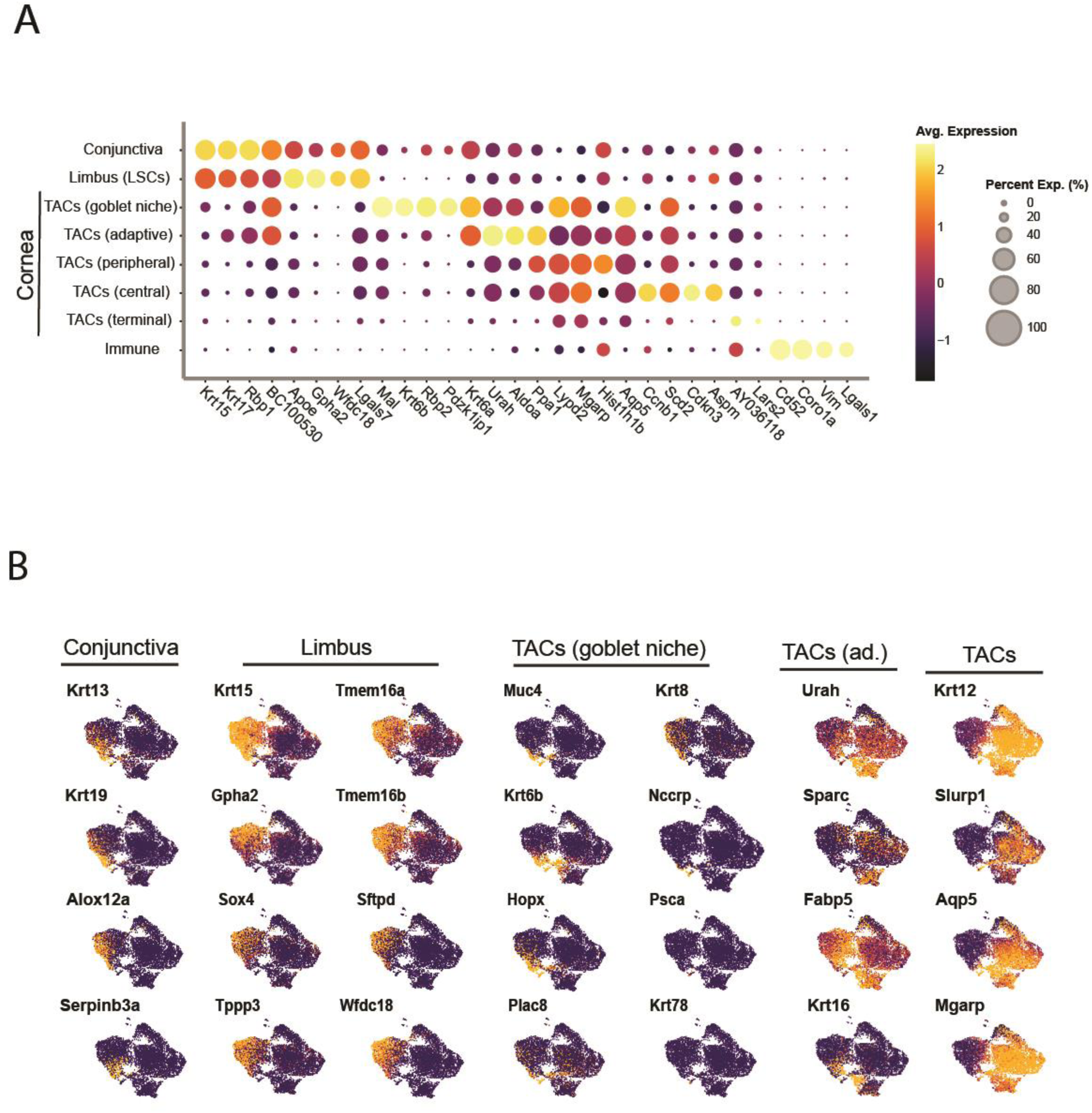
(A) Dot plot of top differentially expressed genes between the identified cell clusters. (B) Umap plots of differentially expressed genes in each of the identified cell clusters.

**Supplementary Figure 3.**
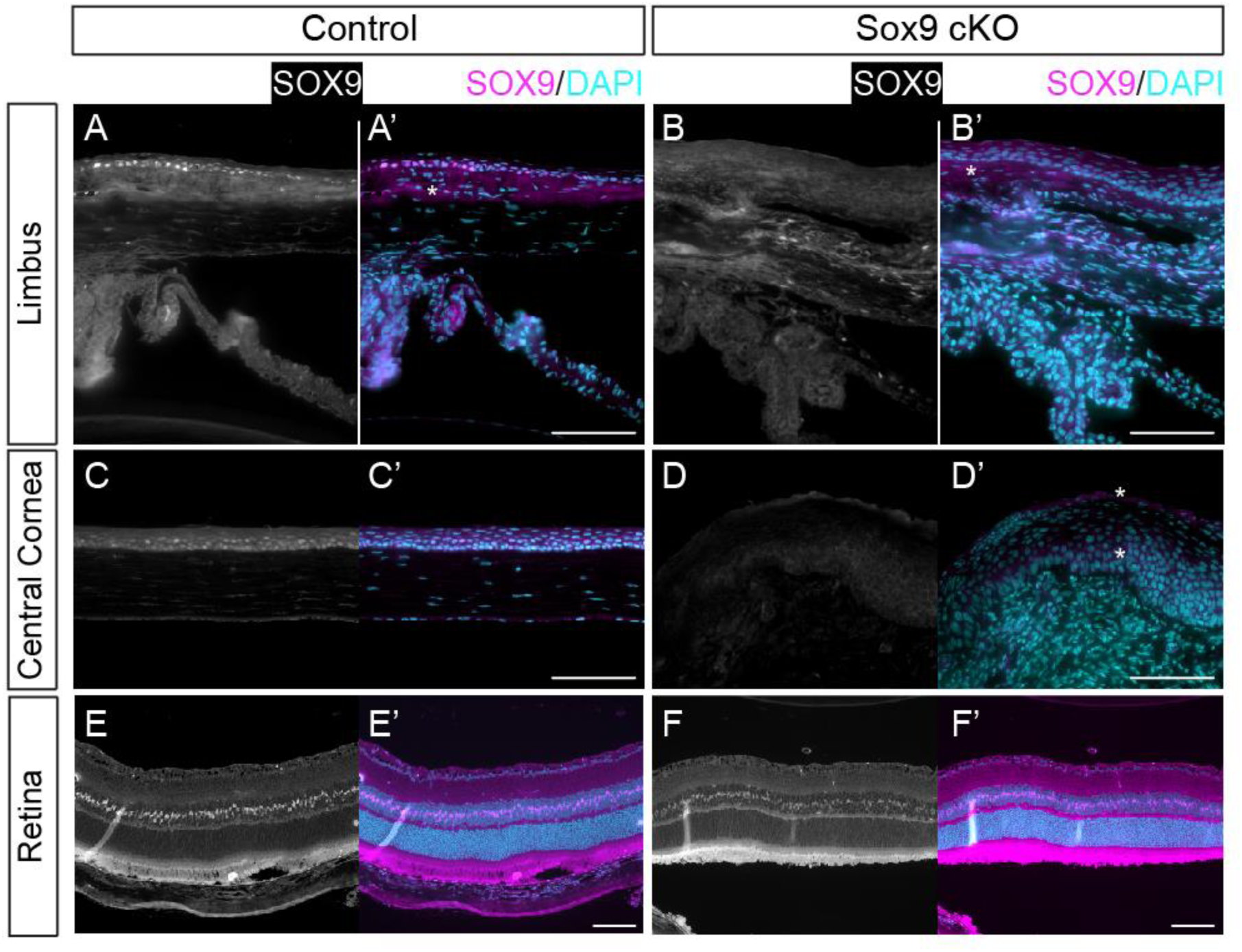
Sox9 Expression in the mouse eye. (A/A’) Immunofluorescence images of Sox9 in the limbus and central cornea (C/C’) of control corneas (n = 25 corneas [13 animals]). (B/B’) Immunofluorescence images of Sox9 in the limbus and central cornea (D/D’) in Sox9-cKO corneas (n = 21 corneas [11 animals]). (E/E’) Immunofluorescence images of control and Sox9-cKO (F/F’) eyes stained for Sox9 in the retina. Muller glial cells in the retina are non-epithelial cells that also express Sox9 but are not targeted for deletion by Cre recombinase. Asterisks indicate non-specific background staining. Scale bars are 100 µm.

**Supplementary Figure 4.**
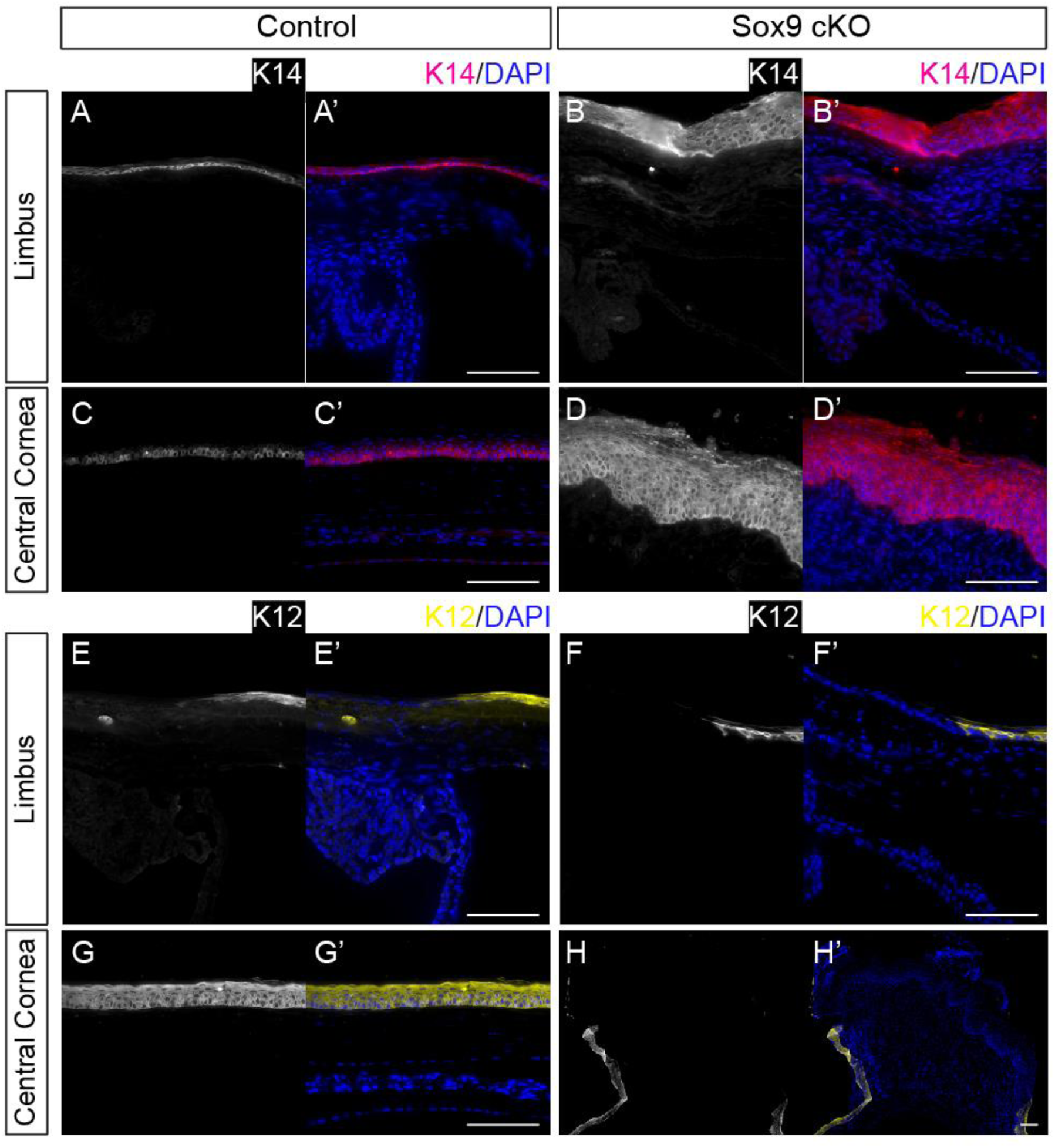
Keratin expression is abnormal in Sox9 cKO corneas. (A/A’) Immunofluorescence images of K14 in the limbus and central cornea (C/C’) of control corneas (n = 25 corneas [13 animals]). (B/B’) Immunofluorescence images of K14 in the limbus and central cornea (D/D’) of Sox9 cKO corneas (n = 21 corneas [11 mice]). (E/E’) Immunofluorescence images of K12 in the limbus and central cornea (G/G’) of control corneas (n = 25 corneas [13 animals]). (F/F’) Immunofluorescence images of K12 in the limbus and central cornea (H/H’) of Sox9-cKO corneas (n = 21 corneas [11 animals]). Keratin 14 (K14), Keratin 12 (K12); Scale bars are 100 µm.

**Supplementary Figure 5.**
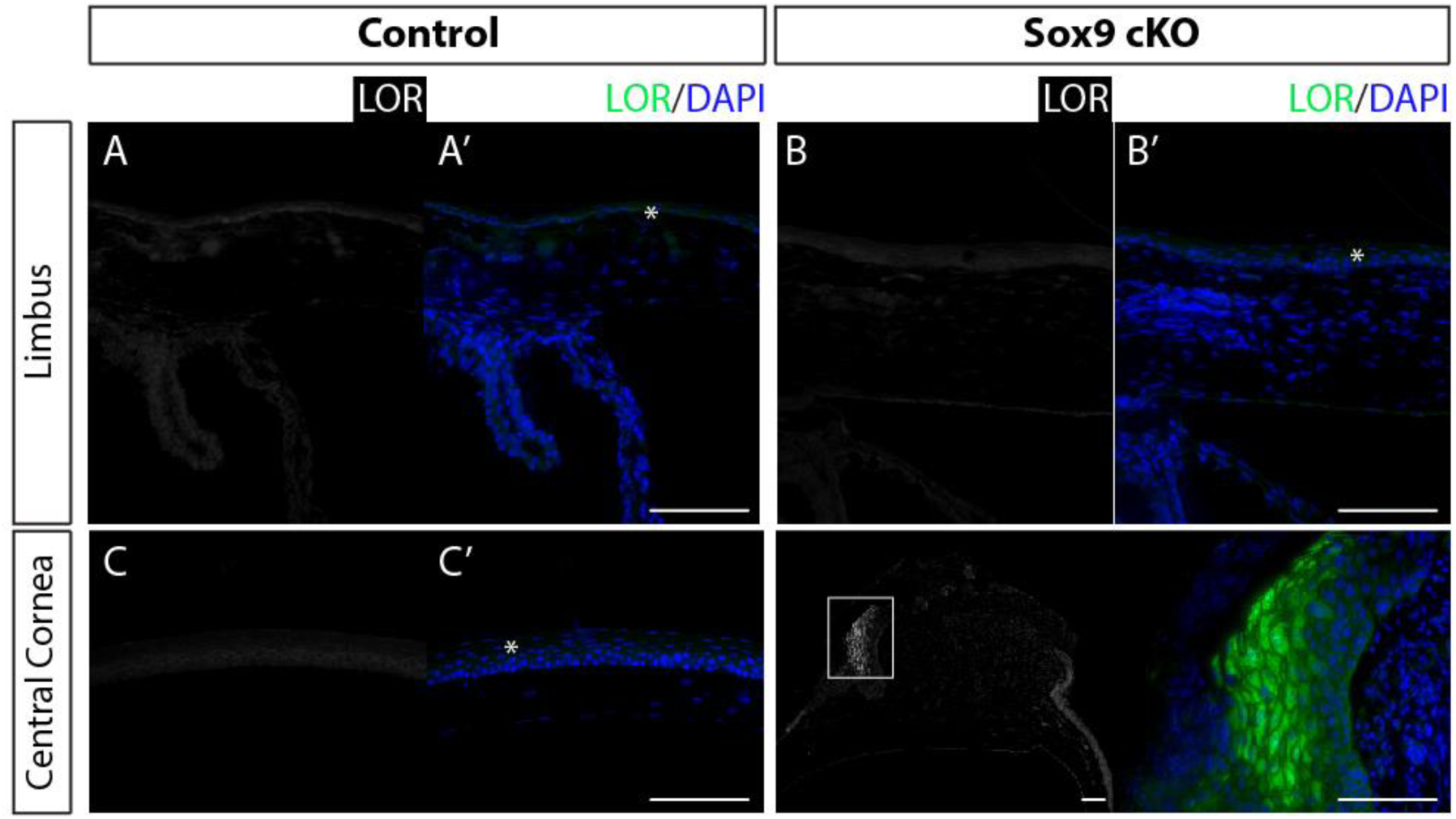
Sox9 cKO corneas express the cornification marker, loricrin. (A/A’) Immunofluorescence images of Loricrin in the limbus and central cornea (C/C’) of control corneas (n = 25 corneas [13 animals]). (B/B’) Immunofluorescence images of Loricrin in the limbus and central cornea (D/D’) in Sox9-cKO corneas (n = 21 corneas [11 animals]). Loricrin (LOR); Asterisks indicate non-specific background staining. Scale bars are 100 µm.

**Supplementary Figure 6.**
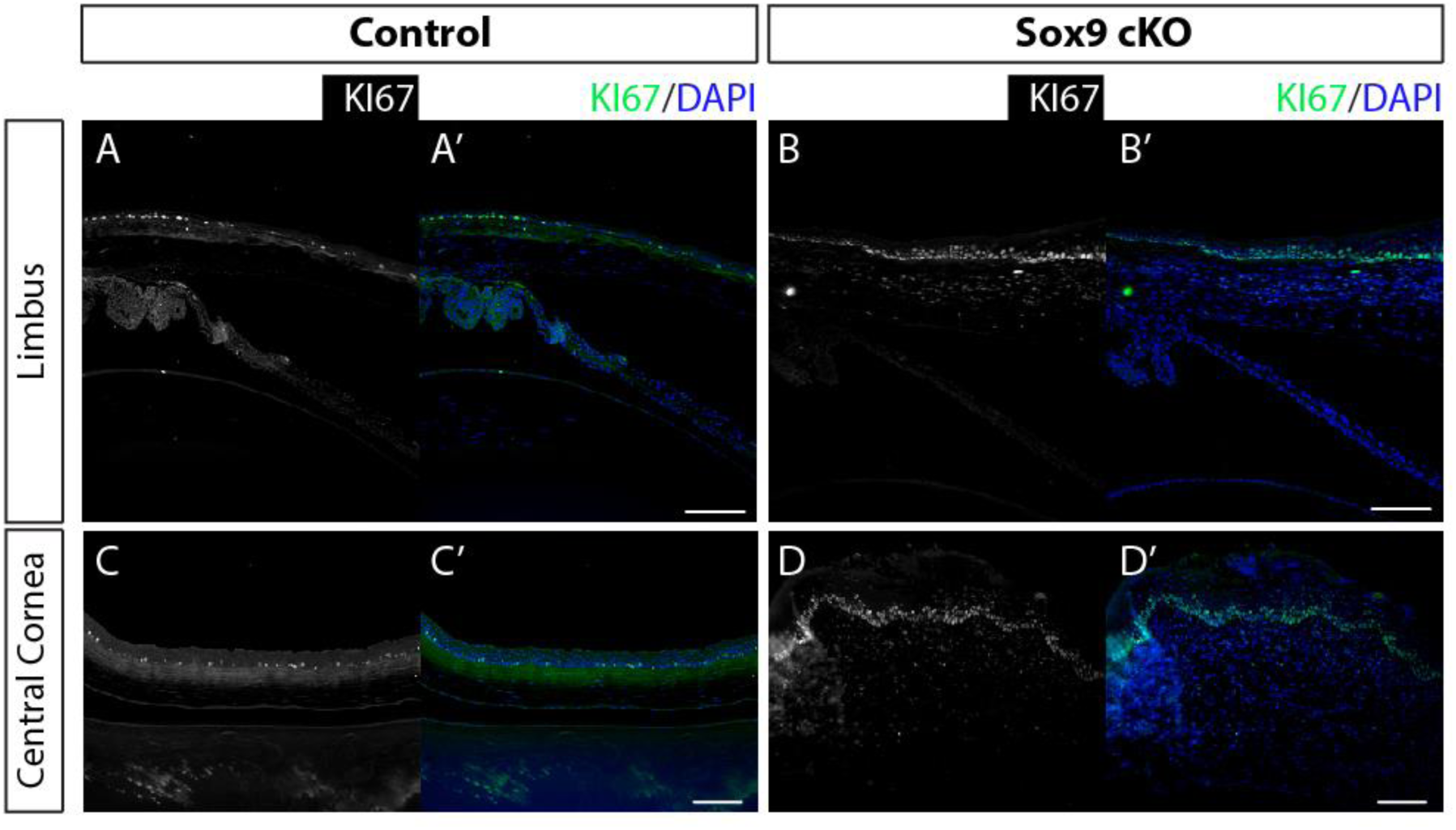
Sox9 loss leads to unchecked proliferation throughout the corneal epithelium. (A/A’) Immunofluorescence images of KI67 in the limbus and central cornea (C/C’) of control corneas (n = 8 corneas [7 mice]). (B/B’) Immunofluorescence images of KI67 in the limbus and central cornea (D/D’) in Sox9-cKO corneas (n = 7 corneas [4 animals]).

